# Development and pre-clinical evaluation of Newcastle disease virus-vectored SARS-CoV-2 intranasal vaccine candidate

**DOI:** 10.1101/2021.03.07.434276

**Authors:** Manolo Fernandez Díaz, Katherine Calderon, Aldo Rojas-Neyra, Vikram N. Vakharia, Ricardo Choque-Guevara, Angela Montalvan, Astrid Poma-Acevedo, Dora Rios-Matos, Andres Agurto-Arteaga, María de Grecia Cauti-Mendoza, Norma Perez-Martinez, Gisela Isasi-Rivas, Luis Tataje-Lavanda, Miryam Palomino, Henri Bailón, Yacory Sernaque-Aguilar, Freddy Ygnacio-Aguirre, Manuel Criollo-Orozco, Edison Huaccachi-Gonzalez, Elmer Delgado-Ccancce, Doris Villanueva-Pérez, Ricardo Montesinos-Millan, Kristel Gutiérrez-Manchay, Katherine Pauyac-Antezana, Ingrid Ramirez-Ortiz, Stefany Quiñones-Garcia, Yudith Cauna-Orocollo, Katherine Vallejos-Sánchez, Angela A. Rios-Angulo, Dennis Núñez-Fernández, Mario I. Salguedo-Bohorquez, Julio Ticona, Manolo Fernández Sánchez, Paquita García, Eliana Icochea, Luis Guevara, Mirko Zimic, for the COVID-19 Working Group in Perú

## Abstract

The COVID-19 pandemic has claimed the lives of millions of people worldwide and threatens to become an endemic problem, therefore the need for as many types of vaccines as possible is of high importance.

Because of the millions of doses required, it is desirable that vaccines are not only safe and effective, but also easy to administer, store, and inexpensive to produce.

Newcastle Disease Virus (NDV) is responsible for a respiratory disease in chickens. It has no pathogenic homologue in humans. NDV is recognized as an oncolytic virus, and its use in humans for oncological treatment is being evaluated.

In the present work, we have developed two types of NDV-vectored candidate vaccines, which carry the surface-exposed RBD and S1 antigens of SARS-CoV-2, respectively. These vaccine candidates were produced in specific-pathogen-free embryonating chicken eggs, and purified from allantoic fluid before lyophilization. These vaccines were administered intranasally to three different animal models: mice, rats and hamsters, and evaluated for safety, toxicity, immunogenicity, stability and efficacy. Efficacy was evaluated in a challenge assay against active SARS-CoV-2 virus in the Golden Syrian hamster model.

The NDV-vectored vaccine based on the S1 antigen was shown to be safe and highly immunogenic, with the ability to neutralize SARS-CoV-2 *in-vitro*, even with an extreme dilution of 1/640. Our results reveal that this vaccine candidate protects the lungs of the animals, preventing cellular damage in this tissue. In addition, this vaccine reduces the viral load in the lungs, suggesting that it may significantly reduce the likelihood of transmission. Being lyophilized, this vaccine candidate is very stable and can be stored for several months at 4-8⁰C.

In conclusion, our NDV-based vaccine candidate has shown a very favorable performance in the pre-clinical study, serving as evidence for a future evaluation in a Phase-I human clinical trial. This candidate represents a promising tool in the fight against COVID-19.

## 1. INTRODUCTION

In December 2019, an outbreak of atypical pneumonia was reported in a market in Wuhan City, China. This was attributed to a virus phylogenetically related to the Severe Acute Respiratory Syndrome Coronavirus (SARS-CoV), which led to an outbreak of severe acute respiratory syndrome in the early 2000s. Later nominated as SARS-CoV-2, this virus is associated with symptoms such as fever, dry cough, and respiratory difficulties. Severe clinical manifestations also include pulmonary and systemic failure with an exacerbated inflammatory condition that can lead to death^1^. The high transmission and mortality, together with the lack of effective treatment, and the high indirect global economic cost, call for the urgent need to develop and evaluate vaccine candidates^2^.

The SARS-CoV-2 virus recognizes the angiotensin-2 converting enzyme (ACE-2), present on the surface of several human cell types, including pneumocytes. The glycosylated Spike (S) protein gives the virus the ability to bind to the cell membrane and promote endocytosis for the entry of the viral particle ^3^. The Spike protein is comprised of two subunits, S1 and S2. The most distal end of the S1 subunit is the receptor binding domain (RBD), which interacts with ACE-2 through the receptor binding motif (RBM)^4^. The SARS-CoV-2 RBD shows structural similarity with its SARS-CoV homolog. However, the structure of SARS-CoV-2 RBM, is associated with higher affinity to ACE-2 than its SARS-CoV counterpart ^5, 6^, giving SARS-CoV-2 a higher virulence ^7–9^.

Previous studies with SARS-CoV and MERS-CoV have aided to identify potential SARS-CoV-2 vaccine candidates, particularly in the use of the S protein due to its known immunogenicity ^10–13^. SARS-CoV-2 has been found to have potential B and T lymphocyte protective epitopes, thus being a promising vaccine candidate^14,15^. An important portion of the S1 surface is glycosylated except for a large epitope in the RBD that mediates the binding to the ACE-2 receptor ^16^. To date, the S1 and RBD are considered important vaccine targets ^17^. The amino acid sequence of S1/RBD is mutating under selection pressure, positively selecting mutants with greater affinity to ACE-2 ^18–22^ and, most recently, mutants that tend to escape neutralization by antibodies against S1 of SARS-CoV-2 ^23,24^. Currently, the development of SARS-CoV-2 vaccines is in full swing, with most vaccine candidates based on the S antigen ^12,25–28^, which has been found to have multiple B and T lymphocyte response sites that would be promising vaccine candidates ^7^.

Different strategies have been applied for the development of vaccines against SARS-CoV-2, seeking safety, effectiveness and protection against the virus, including vaccines based on inactivated virus vaccines based on mRNA, and vaccines using viral vectors ^2,29–31^. The Newcastle disease virus (NDV), is the causative agent of the Newcastle disease (ND), and has been used as a viral vector for the expression of diverse antigens from animal and human pathogens^32–34^. NDV is a member of the *Paramyxoviridae* family, recently known as *Avian orthoavulavirus 1*. NDV is a single-stranded, negative-sense RNA virus with a genome size of approximately 15.2 kb ^35–38^. NDV encodes six structural proteins: nucleocapsid protein (NP), phosphoprotein (P), matrix protein (M), fusion protein (F), haemagglutinin–neuraminidase protein (HN), and the large protein (L), which is a viral polymerase ^39, 40^. The viral particle of NDV has the F and HN glycoproteins, both localized in the envelope of the virion. They participate in the virus attachment, entry to the cell, initiation of the infectious cycle, and release from the cell. NDV can be divided into three groups according to their virulence in poultry: velogenic, mesogenic, and lentogenic ^41^. NDV strain LaSota is lentogenic, and it is routinely used as live NDV vaccine. It grows to a high titer in embryonated chicken eggs, induce strong humoral and cellular immunity and can be administered as drops through the nasal route ^42^.

It has been shown that NDV does not pose a threat to human health, and the majority of the human population have no pre-existing immunity ^33, 42, 43^. NDV has selectivity for tumors, acting as an oncolytic virus. Tumoral cell defects, including anti-viral and apoptotic pathways ^44^, explain the NDV-mediated oncolytic efficiency in mammal cells, through manipulation of antiviral cellular pathways, induction of apoptosis, and indirect activation of the innate and adaptive immune response (humoral, cellular and mucosal) ^32, 45, 46^. The use of oncolytic viruses for cancer treatment in humans have not shown serious side effects unlike other systemic cancer therapies, in extreme cases there could cause mild fever and conjunctivitis ^44^. Important advantages of using NDV vectors are their low (undetectable) rate of recombination and robust production, among others ^47^. For these reasons, some NDV strains are being currently used in human clinical trials for cancer treatment ^45, 48^. NDV has been used as a vector for vaccine development since the late 1990s. The efficiency of vaccines based on this vector has been demonstrated against respiratory viruses, in chickens against infectious bronchitis virus, in monkeys against SARS-CoV and in camels against MERS-CoV ^42, 49–51^.

Previous studies using a NDV-like vector for SARS-CoV and MERS vaccine candidates have provided strong immunogenicity and protection in mice and non-humans primates ^42, 51, 52^. Recently, NDV has been proposed as a potential vector for a vaccine against SARS-CoV-2. Previous studies have shown that it is feasible to produce S protein from other coronaviruses in NDV ^42, 51^. Sun W, *et al.* demonstrated in vivo that a NDV-vectored vaccine against SARS-CoV-2 administered by the intramuscular route, induces a high immune response in mice and hamsters, including some weight loss and reduced viral load in the lungs of challenged animals ^53, 54^. Noteworthy, NDV-vectored vaccines induce mucosal immune response at the respiratory tract, and do not recombine with host DNA during replication ^42^.

In this study, we describe the design and preclinical evaluation of an intranasal NDV-vectored vaccine candidate for COVID-19. The S1 and RBD of the SARS-CoV-2 Spike protein were expressed independently on the surface of a recombinant NDV. These vaccine candidates were tested both in-vitro and in-vivo. Our results show that these constructs are promising candidates in the fight against COVID-19.

## 2. MATERIALS AND METHODS

### 2.1. Ethics statements

Animal research was reviewed and conducted under an approved protocol by the Committee on the Ethics of Animal Experiments of the School of Veterinary and Animal Husbandry of the Universidad Nacional Hermilio Valdizan, Huanuco, Peru. This study was carried out in strict accordance with the recommendations described in the Guide of biosafety and the Care and Use Animals of the National Institute of Health (INS), Lima, Peru ^55^.

### 2.2. Animals

One hundred male and female 4-5 weeks-old Golden Syrian hamsters (*Mesocricetus auratus*) that were, 14 female 5-8 weeks-old albino mice (*Mus musculus*) strain BALB/c, and 36 male 7-9 weeks-old rats (*Rattus norvegicus* albinus, strain Holtzman) were obtained from the Peruvian National Institute of Health (INS).

For the in-vivo assay, all hamsters were transferred and acclimatized to Animal Biosafety Level 3 (BSL-3) facility for 1 week. Then, they were infected with active SARS-CoV-2.

### 2.3. Development and characterization of recombinant NDV expressing SARS-CoV-2 RBD and S1 antigens

#### 2.3.1. Cell Culture

Cells from African green monkey kidney, clone E6 (Vero E6, ATCC^®^ CRL-1586^TM^) and cells derived from Chicken Fibroblast (DF-1, ATCC) (Old Town Manassas, VA, USA), were maintained in Dulbecco’s modified Eagle’s medium (DMEM), supplemented with 5% heat-inactivated fetal bovine serum (FBS), and 1x antibiotic antimycotic (Thermo Fisher Scientific, Waltham, MA, USA). Vero cells (Vero 81, ATCC^®^ CCL-81^TM^) were grown in Eagle’s Minimum Essential Medium (EMEM) supplemented with 10% FBS, 100 IU/mL of penicillin, and 100 µg/mL streptomycin. All cell lines were cultivated at 37°C in an incubator with stable humidity, supplied with 5% CO_2_.

#### 2.3.2. Plasmid Construction

Our vaccine candidates are based on the recombinant lentogenic NDV strain LaSota, which was designed as rLS1 virus. The design and construction of a pFLC-LS1 plasmid (19,319 nucleotides) containing the full-length genome of an infectious NDV clone, and the three support plasmids containing the N, P, and L genes (pCI-N, pCI-P, and pCI-L, respectively) have been previously described ^56^. This NDV-based system is protected under patent 001179-2014/DIN.

The genetic sequences of the RBD and the S1 subunit of the S protein correspond to the SARS-CoV-2 strain isolate from China (GenBank accession no. MN908947.3). To improve the incorporation of RBD and S1 into the NDV virion, we designed two cassettes; 1) the HN-RBD transcriptional cassette (1,013 nt), containing the genetic sequences of the RBD (636 nt), the complete transmembrane domain (TM), and the cytoplasmic tail (CT) of the gene haemagglutinin–neuraminidase (HN) of NDV; and 2) the S1-F transcriptional cassette (2,441 nt), for which the genetic sequence of the S1 subunit (2,043 nt), taken from the S gene (3,822 nt). This sequence was modified to create an open reading frame (ORF), adding the “ATG” nucleotides. Then, fused with the TM and CT of the fusion (F) gene. Both the TM and CT from the HN and F genes were taken from the pFLC-LS1 plasmid. Both transcriptional cassettes were flanked with specific gene-end (GE) and gene-start (GS) transcriptional signals of the paramyxovirus genome ^57^. Further, these cassettes flanked with restriction sites of the *BbvCI* enzyme were chemically synthesized and subsequently cloned into the plasmid pUC57 by GenScript (Piscataway, NJ, USA). These plasmids were purified and DNA extracted using QIAGEN Plasmid Midi Kit (100) (cat. n° 12145, QIAGEN, Valencia, CA, USA), according to manufacturer’s instructions.

The pFLC-LS1 plasmid was digested by the enzyme of unique cut, *BbvCI* (cat. n° R0601L, NEB, New England Biolabs, Ipswich, MA, USA), to obtain the linearized plasmid. Both HN-RBD and S1-F transcriptional cassettes were digested with the same enzyme and then inserted into the P/M junction of the pFLC-LS1 to be expressed as a separate mRNA. The resulting plasmids were designated as pFLC-LS1-HN-RBD (20,315 nt) and pFLC-LS1-S1-F (21,743 nt).

#### 2.3.3. Recovery of the rLS1-HN-RBD and rLS1-S1-F virus

Briefly, the rLS1-HN-RBD and rLS1-S1-F viruses were recovered by co-transfection with a full-length plasmid complementary DNA (cDNA) of each constructs, pFLC-LS1-S1-F and pFLC-LS1-HN-RBD, respectively, together with three support plasmids, as described previously ^56^. The recovered viruses were injected into the allantoic cavities of 9 days-old Specific Pathogen-Free (SPF) embryonated chicken eggs (Charles River Avian Vaccine Services, Norwich, CT, USA). After four days of incubation for at 37°C, the allantoic fluids (AFs) containing the recovered virus were harvested, clarified, aliquoted and stored at −80°C. Presence and recovery of the viruses was detected and confirmed by hemagglutination (HA) assays using 1% chicken red blood cells (RBC). Identification of the recombinant viruses was confirmed by Reverse Transcription-Polymerase Chain reaction (RT-PCR) and by Sanger sequencing (Macrogen Inc., Korea) (data not shown).

#### 2.3.4. Indirect Immunofluorescence Assay (IFA)

To examine the expression of SARS-CoV-2 S RBD and S1 subunit proteins by rLS1-HN-RBD, and rLS1-S1-F, recombinant viruses were developed. Vero-E6 cells were infected with the recombinant viruses and rLS1 at a multiplicity of infection (MOI) of 0.5. After 48 hours post-infection (hpi), the cells were fixed with 4% paraformaldehyde for 25 minutes (min), and then the monolayer was washed three times with Dulbecco’s phosphate-buffered saline (DPBS) and permeabilized with Triton 0.1% X-100 for 15 min at room temperature (RT). Then cells were washed three times with DPBS, and incubated the monolayer with the rabbit polyclonal antibody specific to SARS-CoV-2 RBD protein (1:200) (cat. n° 40592-T62, Sino Biological, Beijing, China), and a chicken antiserum specific to Newcastle disease virus (1:200) (10100482, Charles River Avian Vaccine Services, Norwich, CT, USA) for 1.5h at RT. Next, the monolayer was incubated with Donkey Anti-Rabbit IgG H&L-Alexa Fluor® 594 (1:250) (cat. n° ab150072, Abcam, Cambridge, MA, USA) and Goat Anti-Chicken IgY H&L-Alexa Fluor® 488 (1:1000) (cat. n° ab150169, Abcam, Cambridge, MA, USA) for 60 min at RT. Finally, the cells were marked with 4′,6-diamidino-2-phenylindole (DAPI) (cat. n° ab104139, Abcam, Cambridge, MA, USA), and incubated for 5 min. The results were observed using an ObserverA1 fluorescence microscope (Carl Zeiss, Germany). Digital images were taken at 400x magnification and processed with the AxioCam MRc5 camera (Carl Zeiss, Germany).

#### 2.3.5. Western Blot Analysis

To evaluate the SARS-CoV-2 RBD and S1 subunit proteins expression by rLS1-HN-RBD, and rLS1-S1-F recombinant viruses, Vero E6 cells were infected with the recombinant viruses and rLS1 at an MOI of 1. At 48 hpi the cells were harvested, lysed, and analyzed by Western blot. Additionally, to verify the incorporation of the RBD and S1 subunit proteins into rLS1-HN-RBD and rLS1-S1-F viruses, viral particles from AF of SPF chicken embryonated eggs infected with the recombinant viruses and rLS1, were concentrated by ultracentrifugation (Ultracentrifuge, Beckman, Coulter) at 18,000 revolutions per minute (rpm) at 4°C, and partially purified on 25% sucrose cushion. Western blot analysis was carried out using partially purified viruses from AF and lysate from infected cells, using a rabbit polyclonal antibody specific to SARS-CoV-2 RBD protein (cat. n° 40592-T62, Sino Biological, Beijing, China) (2/5000) as the primary antibody and anti-Rabbit IgG conjugated to HRP (cat. n° A01827, GenScript, Piscataway, NJ, USA) (2/5000) as a secondary antibody. The protein expression was visualized with a CCD camera Azure c600 imaging system (Azure Biosystems, Dublin, USA).

#### 2.3.6. Detection of RBD and S1 subunit proteins on the viral surface by flow cytometry

To determine the presence of RBD on the viral surface of rLS1-HN-RBD, and the presence of the S1 subunit on rLS1-S1-F viruses, virion particles were purified with a 25% sucrose cushion. Vero E6 cells were harvested and washed with DPBS with 5% FBS. 1×10^6^ cells were blocked with DPBS with 5% of normal mouse serum (cat. n° ab7486, Abcam, Cambridge, MA, USA) for 30 min at 37°C. Then, the cells were incubated with rLS1 (0.36 mg/mL), rLS1-S1-F (0.09 mg/mL) or rLS1-HN-RBD (0.2 mg/mL) purified viruses for 30 min at 37⁰C. To remove the residual viral particles not attached to the Vero E6 cells, we washed twice with DPBS and 5% FBS. When this was done the mix was marked with rabbit monoclonal antibody anti-SARS-CoV-2 S1 (1:200) (cat. n° 40150-R007, Sino Biological, Beijing, China) as primary antibody for 1h at 37°C, followed by goat anti-rabbit IgG Alexa Fluor® 488 (1:200) (cat. n°ab150081, Abcam, Cambridge, MA, USA) as secondary antibody. Finally, the cells were analyzed in the flow cytometer FACS Canto II (BD Biosciences, USA). The data obtained was analyzed using the software FlowJo v.10.6 (BD Biosciences, USA), where the percentage of positive cells indicates detection of the SARS-CoV-2 S1 subunit or RBD on the viral surface of viruses bound to Vero E6.

#### 2.3.7. Detection of RBD and S1subunit genes by RT-PCR

To detect the rLS1-HN-RBD and rLS1-S1-F recombinant viruses, viral RNA was extracted from AFs stocks using the QIAamp MinElute Virus Spin kit (cat. n° 57704, Qiagen, Germany), according to manufacturer’s instructions. cDNA was generated from RNA using 4 µL of 5X ProtoScript II buffer (cat. n° M0368L, NEB, New England Biolabs, USA), 2 µL of 0.1 M Dithiothreitol (cat. n° M0368L, NEB, New England Biolabs, USA), 1 µL of 10 mM dNTP (cat. n° N0447L, NEB, New England Biolabs, USA), 0.2 µL of RNase Inhibitor 40 U / µL (cat. n° M0314L, NEB, New England Biolabs, USA), 1 µL of ProtoScript II reverse transcriptase 200 U / µL (cat. n° M0314L NEB, New England Biolabs, USA), 2 µL of random primer mix 60 µM (cat. n° S1330S, NEB, New England Biolabs, USA) and 4.8 µL of nuclease-free water. The cDNA was used as PCR template, using the high-fidelity Master mix Q5 (cat. n° M0492S, NEB, New England Biolabs, USA), and the primers NDV-3LS1-2020-F1 (5’-GATCATGTCACGCCCAATGC-3’) and NDV-3LS1-2020-R1 (5’-GCATCGCAGCGGAAAGTAAC-3’) to amplify the complete insert. The thermal cycling protocol comprised an initial denaturation step at 98°C for 30 seconds (s), followed by 35 cycles of 98°C for 10 s, 72°C for 20 s, 72°C for 30 s for the detection of rLS1-HN-RBD, and 40 s for the detection of rLS1-S1-F. The final extension was carried out at 72°C for 2 min. The PCR products were analyzed by electrophoresis on 1% agarose gels and visualized with a CCD camera Azure c600 imaging system (Azure Biosystems, Dublin, USA).

#### 2.3.8. Genetic stability of the rLS1-HN-RBD and rLS1-S1-F virus

The genetic stability of the recombinant viruses across multiples passages was evaluated on 9 days old-SPF embryonated chicken eggs, Viral RNA was extracted from purified viruses of the 3^rd^ and 6^th^ passage, where the presence of the gene inserts was confirmed by RT-PCR using specific primers. The expression of the SARS-CoV-2 S1 subunit and RBD inserts were also evaluated using purified viruses of the 3^rd^ and 6^th^ passage by Western blotting.

#### 2.3.9. In vitro replication properties of the rLS1-HN-RBD and rLS1-S1-F viruses, plaque assay, and pathogenicity

We compared the infectivity and growth properties between the rLS1-HN-RBD, rLS1-S1-F, and rLS1 viruses. DF-1 cells were seeded in monolayer culture at 70% confluence in 12-well plates and infected with rLS1-HN-RBD, rLS1-S1-F, and rLS1 viruses at MOI 0.05. Cells were maintained with DMEM containing 1% FBS and 5% AF and incubated at 37°C with 5% CO_2_. Supernatants of the infected cells were collected at 12, 24, 36, 48, 60, and 72 hpi and kept at −80°C. Titers of each collected supernatant were determined using plaque assay, as previously described ^56^. These experiments were repeated at 3 specific time points. In addition, the morphology and size of the plaques of the two recombinant viruses were compared with those formed with rLS1 infection. To determine the pathogenicity, the viruses were evaluated in Mean Death Time (MDT) and Intracerebral Pathogenicity Index (IPIC) assays in 10 days-old SPF embryonated chicken eggs and one-day-old SPF chickens (Charles River Avian Vaccine Services, Norwich, CT, USA), respectively, using standard procedures ^58^.

#### 2.3.10. Preparation and stability of the lyophilized vaccine

As preparation for a future production system, the rLS1-RBD-HN and rLS1-S1-F viruses were separately inoculated into allantoic cavities of 9 to 11 days-old SPF embryonated chicken eggs. After four days of incubation at 37°C, the AFs were harvested, clarified, and filtered using 0.22 µm filters. The presence of the viruses in AFs was detected and confirmed by HA. Finally, the AFs containing the rLS1-RBD-HN, rLS1-S1-F, and the mixture of both viruses were placed in vials (2 mL/vial) and lyophilized using an MX5356 lyophilizer (Millrock Technology). The lyophilized vaccine of the mixture of rLS1-HN-RBD and rLS1-S1-F viruses were stored at 4°C and were evaluated by plaque assay, HA, and Western blot assays on days 1, 30, and 50 after lyophilization. The lyophilized vaccines were used in the following in vivo tests in the animals of the study.

### 2.4. Safety-like assay in mice

Eighteen mice weighing between 23-27 g were immunized intranasally (i.n.) on days 0 and 15 (prime-boost respectively), with 5×10^6^ PFU/mice (40µL volume) of rLS1-HN-RBD (*n* = 2), rLS1-S1-F (*n* = 4), rLS1-HN-RBD/rLS1-S1-F (*n* = 4). We reserved an unvaccinated group (*n* = 4) as control. Thirty days after the boost (day 45), all animals were anesthetized with 100 µL of Ketamine (100 mg), Xylazine (20 mg), and Atropine Sulfate (1 mg) via intramuscular (i.m.) injection and sacrificed. The lungs were collected for histopathological analysis, which was performed in an Axio Scope.A1 microscope (Zeiss, Germany).

### 2.5. Toxicity-like assay in rats

Forty-two rats weighing between 340-360 g were used. Six groups of rats (*n*=7) were immunized i.n. with 40 µL of the rLS1-S1-F vaccine on days 0, 7, and 14, group 1 (1.39×10^7^ PFU/kg), group 2 (3.49×10^7^ PFU/kg), group 3 (8.76×10^7^ PFU/kg), group 4 (2.20×10^8^ PFU/kg, group 5 (5.53×10^8^ PFU/kg), group 6 (1.39×10^9^ PFU/kg) and the control group received just only allantoic fluid. They were monitored for 21 days from the start of the experiment, and the mortality and body weight variation were documented weekly.

### 2.6. Immunogenicity in hamsters

Forty-eight Golden Syrian hamsters weighing between 120-140 g were divided into 4 groups (*n* =12 per group): group 1 (rLS1-HN-RBD), group 2 (rLS1-S1-F), group 3 (rLS1-HN-RBD/rLS1-S1-F), and the unvaccinated group 4 (control). They were immunized i.n. with 5×10^6^ PFU/hamster (40 µL volume) following a prime-boost regimen with a two-week interval. Immunized hamsters were bled right before the boost and fifteen days post-boost (at days 15 and 30 respectively), to measure the SARS-CoV-2 RBD and S specific serum IgG antibody by indirect ELISA assay, as well as the neutralizing antibody (nAbs) titers using a surrogate Virus Neutralization Test (sVNT) and by Plaque Reduction Neutralization Test (PRNT) against SARS-CoV-2 virus.

#### 2.6.1. Enzyme-Linked Immunosorbent Assay (ELISA) indirect IgG

Immunized hamsters were bled on days 15 and 30 after immunization. Serum from each sample was isolated by centrifugation at 2500 rpm for 5 min. To perform the assay, Nunc MaxiSorp 96-well flat-bottom plates (cat. n° M9410 Sigma-Aldrich) were coated with 100 µL of SARS-CoV-2 RBD (1 µg/mL) (cat n° Z03479, GenScript, Piscataway, NJ, USA) and S1 subunit (0.5 µg/mL) (cat n° Z03501, GenScript, Piscataway, NJ, USA) purified recombinant proteins dissolved in carbonate-bicarbonate buffer (pH 9.6) and incubated at 4°C overnight. The next day, the wells were washed six times with DPBS containing 0.05% (v/v) Tween-20 (PBS-T 0.05%) and blocked with 3% (w/v) Difco Skim Milk (cat. n° 232100, BD Biosciences, USA) in PBS-T 0.05% for 2 h in agitation at RT. The plates were then washed six times with DPBS-T 0.05%. Then, 100 µL of each collected serum sample diluted 1:100 with 1% (w/v) Difco Skim Milk (cat. n° 232100, BD Biosciences, USA) was added to each plate for 1 h at 37°C. The wells were washed six times with PBS-T 0.05% and further incubated with 100 µL (1:28000) of Goat Anti-Syrian Hamster IgG conjugated to HRP (cat. n° ab6892, Abcam, Cambridge, MA, USA) diluted in 1% Difco Skim Milk in PBS-T 0.05% for 1h at 37°C. The plates were washed six times with PBS-T 0.05% and they were incubated with 100 µL of 3,3’,5,5’-tetramethylbenzidine (TMB) on the plate for 15 min at RT. Finally, the reaction was stopped by adding 50 µL per well of 2 N H_2_SO_4_, and the plates were read at 450 nm using an Epoch 2 microplate reader (Biotek, USA). The negative control was obtained from serum samples of the control group.

#### 2.6.2 Neutralization Tests using SARS-CoV-2 surrogate virus

Serum samples were processed to evaluate nAbs titers against SARS-CoV-2. All neutralization assays performed with the surrogate Virus Neutralization Test (sVNT) (cat. n° L00847, GenScript, Piscataway, NJ, USA) following manufacturer’s instructions. Plates were read using an Epoch 2 microplate reader (Biotek, USA) at 450 nm. The positive and negative cut-offs for SARS-CoV-2 nAbs detection were interpreted as inhibition rate, as follows: *positive*, if ≥ 20% (neutralizing antibody detected), and *negative*, if <20% (neutralizing antibody no detectable).

#### 2.6.3. Plaque Reduction Neutralization Test (PRNT) of SARS-CoV-2 virus Isolation of SARS-CoV-2

SARS-CoV-2 (28549) was isolated from a nasopharyngeal swab sample collected from a patient with confirmed RT-PCR diagnostic for SARS-COV-2 infection in April of 2020 in Lima, Peru by the INS. The identity of the virus was confirmed by whole genome sequencing. Virus isolation was performed in Vero cells ATCC ® CCL 81 TM in Eaglés Minimum Essential Medium (EMEM) medium supplemented with 10% fetal bovine serum (FBS), 100 IU /mL of penicillin and 100 µg/mL streptomycin and cultured at 37 °C in an incubator with humid atmosphere at 5% CO_2_. The sample was filtered through 0.22 μm pore membrane and inoculated with 100 μL into a confluent monolayer of Vero 81 cell line. Cells were observed daily to detect the appearance of any cytopathic effect and collected it for confirmation. The virus was propagated in Vero 81 cell culture for viral stock production at −80⁰C and titer determined by PFU.

##### Plaque Reduction Neutralization Test

Vero E6 cells were seeded in 24-well plates (Corning Costar, NY, USA) at a confluence of 90%. Pooled hamster serum samples were collected at day 30 of immunization. They were heat-inactivated (HI) at 56°C for 30 min, then after two-fold serial dilutions were mixed and incubated with 40-50 PFUs of SARS-CoV-2 (28549) for 1 h at 37°C in 5% CO_2_. The serum-SARS-CoV-2 mixtures were added to 24-well plates of Vero E6 and incubated at 37°C and 5% CO_2_ for 1 h. After absorption, the serum-virus mixtures were removed, and a liquid overlay medium (L-OM) comprising 0.75% carboxymethylcellulose (CMC) (cat. n° C4888-500G, Sigma-Aldrich) supplement with 2% FBS was added to the monolayer cells, that were incubated at 37°C and 5% CO_2_ for 5 days. The plates were fixed and stained with 10 % formaldehyde and 0.5% crystal violet solution ^59^. Each serum was tested in duplicates. The plates were enumerated counted for the calculation of PRNT_50_, considered the gold standard method ^60^.

#### 2.6.4. Cellular immunity for Cytokines quantification by qPCR

Spleens were obtained from hamsters immunized with rLS1-HN-RBD, rLS1-S1-F, and rLS1-HN-RBD/rLS1-S1-F and AF (MOCK) 15 days post-immunization, and transported in RNAlater (cat. n° AM7021, Thermo Fisher Scientific, USA), stored at 4°C overnight and then at –80°C.

RNA was extracted with RNeasy Mini kit (cat. n° 74106, Qiagen, Hilden, Germany), treated with the TURBO DNA-free™ kit (Thermo Fisher Scientific, USA), and converted to cDNA with ProtoScript® II First Strand cDNA Synthesis Kit (cat. n° E6560L, NEB, New England Biolabs, Ipswich, MA, USA) according to manufacturing specifications. cDNA was stored at –20°C until analysis.

Cytokines interferon-gamma (IFNγ), Tumor Necrosis Factor-Alpha (TNF-α) and interleukin-10 (IL-10), and reference gene β-actin were evaluated with primers previously reported ^61–64^. Standard curves were made for all primers, obtaining acceptable efficiency and R^2^ values (data not shown). Master mix preparation and cycling conditions were realized with Luna® Universal qPCR Master Mix kit (cat. n° M3003E, NEB, New England Biolabs, Ipswich, MA, USA) according to manufacturer’s instructions. Briefly, we used 5 µL of the sample (∼ 2 ng/µL cDNA) with 2-3 technical replicas. The qPCR experiments were done on the Rotor-Gene Q equipment (Qiagen, Hilden, Germany) and the ΔΔCT method ^65^ was used for data analysis.

#### 2.6.5. Cellular immunity for Cytokines quantification by ELISA

Whole blood was obtained from hamsters immunized with rLS1-HN-RBD, rLS1-S1-F, and rLS1-HN-RBD/rLS1-S1-F and AF (MOCK) at 15 days post-immunization, was centrifuged at 1000 x g for 20 min at 4°C to obtain the serum. This was duly aliquoted, frozen, and stored at −80°C until analysis. The following quantitative ELISA kits wereused: hamster TNFα ELISA Kit (MBS1600809, San Diego, CA, USA); hamster Interleukin-2 (IL-2) ELISA Kit (MBS033947, San Diego, CA, USA); hamster IFN-γ ELISA Kit (MBS7606840, San Diego, CA, USA); hamster interleukin-4 (IL4) ELISA Kit (MBS706646, San Diego, CA, USA), and hamster IL10 ELISA Kit (MBS706850, San Diego, CA, USA), all procedures were performed following the manufacturer’s instructions. Briefly, the sera were placed 96-well plates in duplicate, and pre-coated with antibodies against the hamster’s cytokines: TNFα, IFNγ, IL-2, IL-4, and IL-10. The plates were incubated at 37 °C and revealed with streptavidin or avidin conjugated with peroxidase (HRP), to finally add the substrate 3,3’, 5,5’-tetramethylbenzidine (TMB), stopping the reaction with sulfuric acid. Plates were read in the EON spectrophotometer (Biotek, USA) at 450 nm. The level of cytokines (pg/mL) detected in the serum of the animals vaccinated with rLS1-HN-RBD, rLS1-S1-F, and rLS1-HN-RBD/rLS1-S1-F were contrasted with MOCK animals.

### 2.7. Efficacy of the vaccines against SARS-CoV-2 challenge

Forty-eight golden Syrian hamsters, divided into 4 groups (*n* =12): group 1 (rLS1-HN-RBD), group 2 (rLS1-S1-F), group 3 (rLS1-HN-RBD/rLS1-S1-F), and the unvaccinated control group 4, these were i.n. challenged with 1 x 10^5^ PFU/hamster in DMEM (40 µL volume) of SARS-CoV-2 at 45 days post-prime immunization.

Four animals in each group were anesthetized and sacrificed with one overdose of 1mL of a mixture of Ketamine (100 mg), Xylazine (20 mg), and Atropine Sulfate (1 mg) by i.m. injection at 2, 5, and 10 days post-challenge. The samples lung tissues (right and left lobes) were separated into two parts: (1) The right lobe was used for the pathological examination, and (2) the left lobe was immediately frozen at −80°C until used; this lobe was used for live infectious virus by viral isolation.

The SARS-CoV-2 (28549) virus specimen used in the challenge, was kindly provided by the National Institute of Health (INS), Lima, Peru. All work and handling with SARS-CoV-2 were performed in a BSL-3 laboratory following the biosafety guidelines of INS.

#### 2.7.1. Histopathology analysis

Lungs were obtained from sacrificed hamsters at days 2, 5, and 10 post-challenge with SARS-CoV-2; and fixed in 10% buffered formalin for 48 h. Organs were then reduced and placed in a container for 24 h with buffered formalin. The containers with the organs were passed to an automatic tissue processor (Microm brand) conducting the following processes: dehydration, diaphanating, rinsing, and impregnation; within an average of 8 h. Organs included in paraffin were cut to a thickness of 5 microns (Microtome Leica RM2245 of disposable metal blades) and placed in a flotation solution in a water bath and then fixed on a slide sheet, dried in the stove (at 37°C for 1 to 2 h). The staining was done with the Hematoxylin and Eosin staining method (H&E), in a battery of staining bottles, as follows: remove paraffin, hydration, hematoxylin coloration, washing, Eosin coloration, rinsing, dehydration, drying, rinsing, then mounting in a microscope slide with Canada Balm (glue), and drying (at 37°C for 12 to 24 h), for further labeling. The final slides colored with H&E were taken and analyzed under an AxioCam MRc5 camera and AxioScope.A1 microscope (Carl Zeiss, Germany) at an amplitude of 20 and 40x by a board-certified veterinary pathologist.

#### 2.7.2. Viral viability: Culture and immunofluorescence assay (IFA)

For virus viability, 60 lung tissue samples from challenged animals were crushed and homogenized in 5 % w/v of EMEM 1% antibiotic antimycotic and centrifuged at 10,000 rpm for 10 min at 4°C. The supernatant was filtered with a 0.22 μm pore Millipore pore filter membrane, then 100 µL inoculated into a confluent monolayer of the Vero 81 cell line, and cultured at 37°C in an incubator with humid atmosphere at 5% CO_2_. The cultures were observed daily for 10 days through the inverted microscope. Lung virus isolation was confirmed by RT-PCR, as described previously ^66^. The IFA was performed using a polyclonal antibody against SARS-CoV-2 from convalescent patients of COVID-19 disease, and anti-human IgG peroxidase conjugate (Sigma).

#### 2.7.3. Animal mobility

To assess hamster’s mobility (in groups 1 to 4) post-challenge, the average velocity, average acceleration, and average displacement were calculated based on videos with a camera positioned on top of the hamsters. Videos of days 2, 5, and 10 post-challenge were analyzed.

Of note, conditions of video recording (distance and focus) were kept constant for consistency. Therefore, the spatial separation between pixels is always the same. Since hamsters do not necessarily move a lot at the border of the box, we estimated average velocity, acceleration and displacement based on any movement occurring away from the box edges. Therefore, movement along the box edges were excluded. We tracked movement throughout 2-3 min time intervals (Supplementary Figure 2). Then, hamsters were tracked in time intervals with no interaction with the box edges. Tracking was carried out using the Kernelized Correlation Filter (KCF)^67^.

This tracking algorithm was developed using the OpenCV library of the Python programming language. Tracking the hamsters produced a video of the X and Y positions of the hamster in the image. Finally, once the tracking record was obtained at the intervals of interest, the average velocity, acceleration, and displacement were calculated for each hamster.

#### 2.7.4. Animal weight variation

The body weight change was measured at days 2, 5, and 10 post-challenge. An additional MOCK group (*n*=12) of unvaccinated not-challenged animals outside the BSL3 were evaluated. These measurements were used to calculate the percentage of body weight variation, comparing to the day 0 of each animal.

### 2.8. Statistical analysis

For the statistical analysis of weight variation in rat groups, we used the one factor analysis of variance (ANOVA). To compare treatments based on the quantification of Cytokines, by qPCR and ELISA, we used the non-parametric Mann-Whitney-Wilcoxon test. To evaluate the statistical significance of body weight change in hamster groups, a one-way ANOVA with multiple comparisons for all the treatments involved was performed. To evaluate changes in hamsters’ mobility over time, non-parametric statistics using the Mann-Whitney and Kruskal-Walls tests were performed using the SciPy v1.5.2 package. To assess plaque reduction (%) of neutralization from the different groups of hamsters, we used two-way ANOVA and Tukey’s post hoc. In all analyses, the statistical software STATA v.16 was used, with a 5% significance level.

## 3. RESULTS

### 3.1. Development and characterization of recombinant NDV expressing SARS-CoV-2 RBD and S1 antigens

#### 3.1.1. Generation of rNDVs expressing RBD and S1subunit genes of SARS-CoV-2

Vero-E6 cells were co-transfected with full-length plasmid cDNA of constructs pFLC-LS1-HN-RBD and pFLC-LS1-S1-F together with three supporting plasmids encoding the NP, P, and L proteins of NDV, essential for replication of NDV (Figure 1A, 1B). At 72 h post-transfection, the cells showed several visible plaques with typical cytopathic effect (CPE) from NDV, demonstrating the successful rescue of both recombinant viruses. The supernatants that were collected five days after transfection were injected into the allantoic cavities of 9-days-old embryonated SPF eggs. The AFs were harvested four days after inoculation and analyzed by HA using chicken RBC. We found positive HA titers ranging from 2 to 2048.

**Figure 1.**
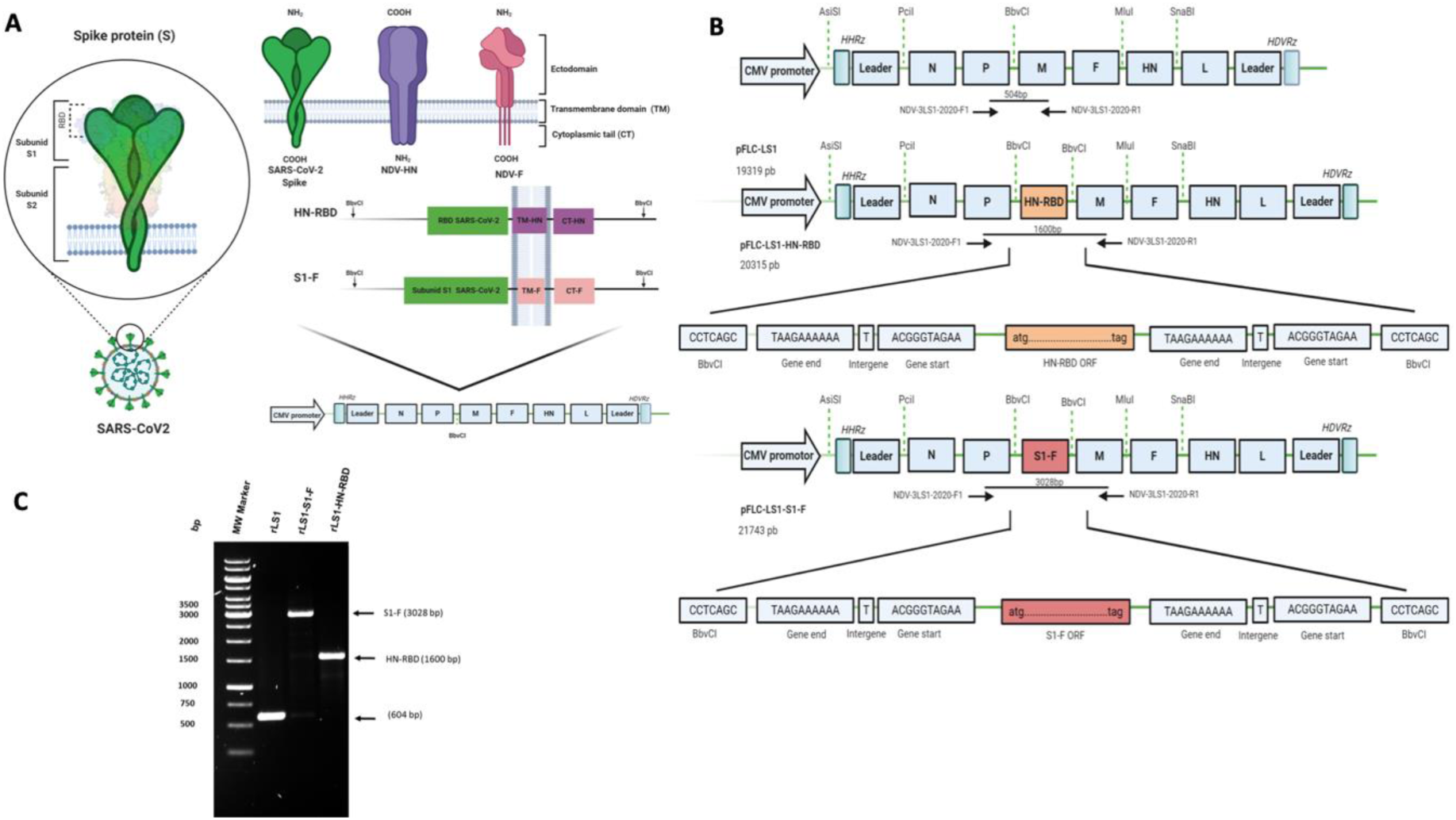
The strategy used for the generation of the recombinant NDVs expressing SARS-CoV-2 RBD and S1. (**A**) The schematic representation of the strategy of construction recombinant NDVs. Two cassettes transcriptional were designed for expressing RBD and S1: 1) HN-RBD was fused with the complete transmembrane domain (TM) and the cytoplasmic tail (CT) of the gene haemagglutinin–neuraminidase (HN), 2) S1-F was fused with the TM/CT of the gene fusion (F) from of full-length pFLC-LS1. (**B**) The full-length antigenome of NDV strain LaSota was used as a clone (pFLC-LS1) was used as back clone, the pFLC-LS1-HN-RBD and pFLC-LS1-S1-F were generated from cassettes that expressing RBD and S1 inserted into genome NDV under control of transcriptional gene end (GE) and gene start (GS) signals. The names, position, and direction of the primers used are shown with arrows (blacks) indicating size products. (**C**) The insertion of the expression cassette into the non-coding region between the P/M genes of NDV genome was verified by RT-PCR using the junction primers NDV-3LS1-2020-F1 and NDV-3LS1-2020-R1 as shown in (B).

The presence of the HN-RBD and S1-F expression cassettes inserted into the non-coding region between the P and M genes of NDV genome was verified by RT-PCR, yielding fragments of 1600 and 3028 bases pairs (bp), which were subsequently amplified and sequenced using junction primers NDV-3LS1-2020-F1 and NDV-3LS1-2020-R1, demonstrating proper insertion into the NDV genome (Figure 1C). The new recombinant NDV viruses were named: rLS1-HN-RBD and rLS1-S1-F.

#### 3.1.2. Expression of the SARS-CoV-2 Proteins in rLS1-HN-RBD and rLS1-S1-F Viruses

Two protein bands with a molecular mass of ∼90 kDa (S1-F) and ∼30 kDa (HN-RBD) were detected in cell lysates infected with rLS1-S1-F and rLS1-HN-RBD viruses respectively (Figure 2A). Two protein bands were detected in the purified recombinant virus (Figure 2B), confirming that the S1-F or HN-RBD proteins were incorporated into the viral particles of rLS1-S1-F and rLS1-HN-RBD viruses respectively. These protein bands were not detected in the rLS1-infected cells and purified viral particles from rLS1 virus.

**Figure 2.**
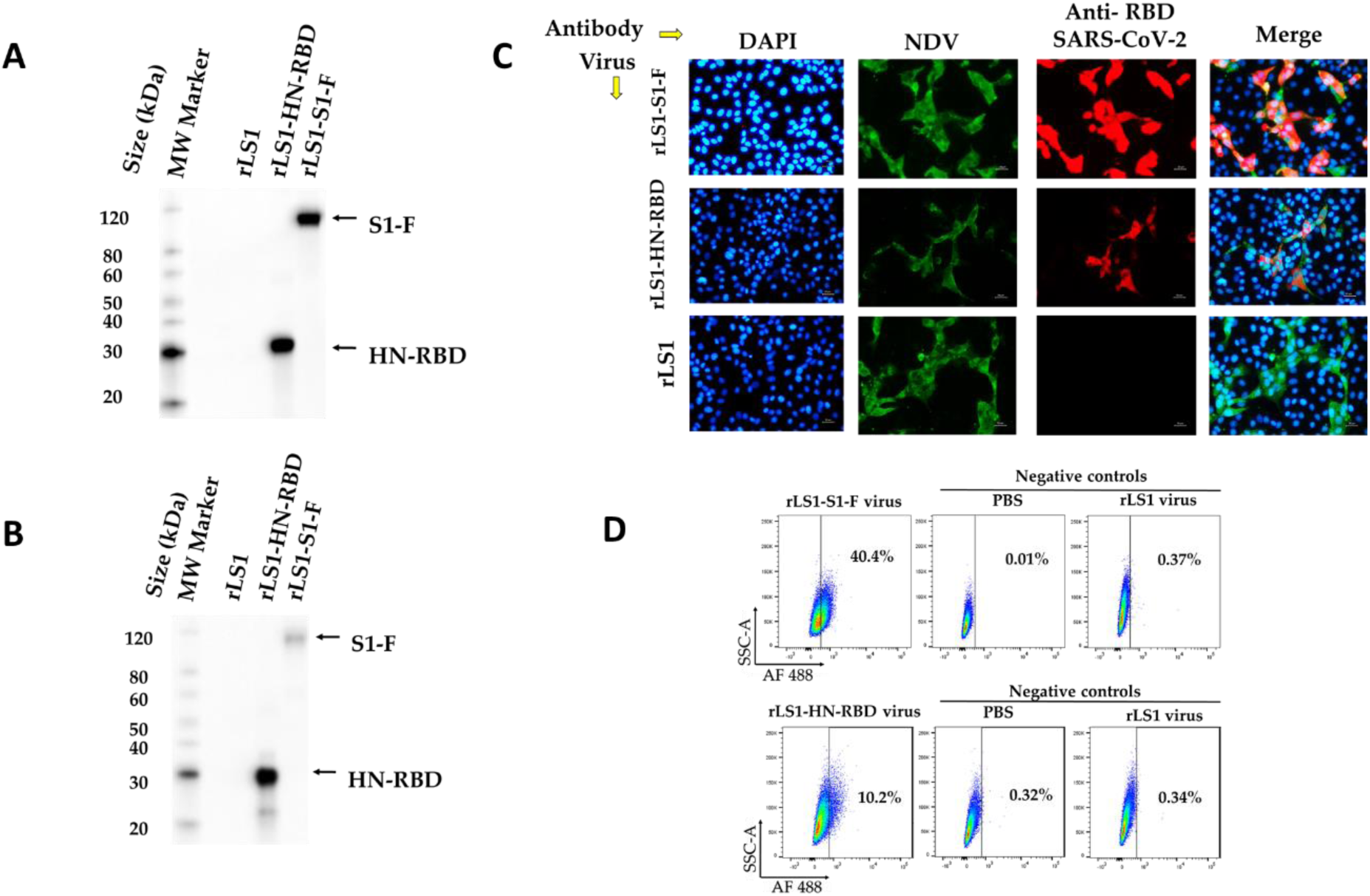
Expression of SARS-CoV-2 RBD and S1 proteins in infected Vero E6 cells and NDV particles. (**A)** Western blot detection for the HN-RBD and S1-F proteins expression. Vero E6 cells were infected with the rLS1, rLS1 rLS1-HN-RBD, and rLS1-S1-F viruses at an MOI of 1. Then after 48 hpi, the cells were lysed and analyzed by western blotting. (**B)** To verify the incorporation of the HN-RBD and S1-F proteins into rLS1-HN-RBD, and rLS1-S1-F viruses, the viral particles in FA from infected SPF chicken embryonated eggs with the recombinant viruses and rLS1, was concentrated by ultracentrifugation, and partially purified on 25 % sucrose cushion. Western blot analysis was carried out using partially purified viruses and lysate from infected cells, using a rabbit antibody specific to SARS-CoV-2 RBD protein and Anti Rabbit IgG conjugated to HRP. The black arrow indicates the expected protein band. (**C**) Vero-E6 cells infected with the rLS1, rLS1-HN-RBD, and rLS1-S1-F at an MOI of 0.5. After 48 h, the expression of RBD and S1 proteins was detected by Immunofluorescence assay using a rabbit antibody specific to SARS-CoV-2 RBD protein, and a Donkey Anti-Rabbit IgG H&L-Alexa Fluor 594. Therefore, the NDV was detected using a chicken antiserum specific to the NDV, and a Goat Anti-Chicken IgY H&L-Alexa Fluor® 488. Cell nuclei were stained with DAPI. A scale bar of 50 µm. Image magnification 200x. (**D**) Detection of S1 or RBD proteins on the viral surface of rLS1-S1-F and rLS1-HN-RBD viruses’ attachment to Vero E6 cells. The cells were incubated with purified viruses rLS1-HN-RBD or rLS1-S1-F, for 30 min. Subsequently, the cells were labeled with rabbit monoclonal antibody anti-SARS-COV-2 S1 as the primary antibody, followed by secondary antibody goat anti-rabbit IgG Alexa Fluor 488. The cells were then analyzed by a flow cytometer. The percentage of positive cells indicates the detection of S1 or RBD proteins on the viral surface of viruses bound to Vero E6 and is shown in the dot plot for rLS1-S1-F virus and sLS1 -HN-RBD virus; including negative controls for each assay determined by cells incubated with phosphate-buffered saline (PBS) or rLS1 virus.

The expression of SARS-CoV-2 RBD and S1 subunit was detected in Vero-E6 cells infected with rLS1-HN-RBD, and rLS1-S1-F by Immunofluorescence assay. RBD and S1 subunit expression was not detected in the cells infected with the rLS1 virus. The NDV protein expression was detected using a chicken antiserum specific to the NDV, and a Goat Anti-Chicken IgY H&L-Alexa Fluor® 488 in Vero-E6 cells infected with rLS1, rLS1-HN-RBD, and rLS1-S1-F viruses (Figure 2C).

Detection of SARS-CoV-2 S1 subunit or RBD on the viral surface of rLS1-S1-F and rLS1-HN-RBD viruses bound to Vero E6 cells was confirmed by flow cytometry (Figure 2D). For the rLS1-S1-F virus, 40.4% positive cells were detected, for the rLS1-HN-RBD virus 10.2% positive cells were detected, and for the rLS1 virus up to 0.37% positive cells were detected. For rLS1-S1-F virus, a higher percentage of cells was detected compared to rLS1-HN-RBD virus.

#### 3.1.3. In Vitro replication properties, genetic stability, and pathogenicity of the rLS1-HN-RBD and rLS1-S1-F viruses

The rLS1-HN-RBD and rLS1-S1-F viruses showed similar growth patterns in DF-1 cells compared to the rLS1 virus (Figure 3A). The virus plaque morphology and sizes of the three viruses in DF-1 cells at 96 hpi were very similar and no notable difference in replication rates between the three viruses (Figure 3B) was observed.

**Figure 3.**
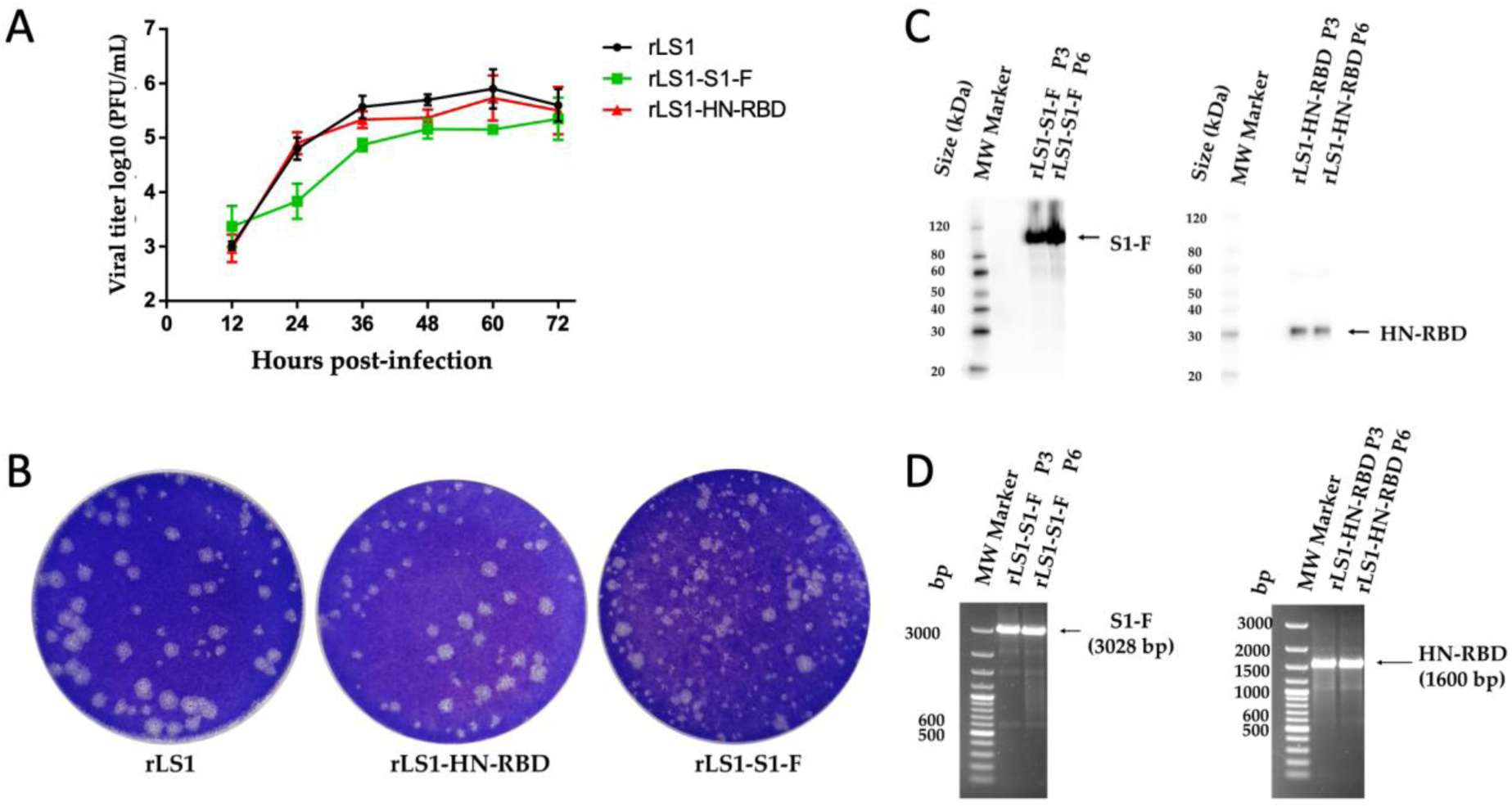
Plaque morphology, replication properties NDVs and genetic stability of the recombinants (A) DF-1 cells were infected with either the rLS1-HN-RBD, rLS1-S1-F, and rLS1 viruses at 0.05 MOI. The infected cells were harvested at 24, 48, and 72 hpi and titrated by plaque assay in triplicates. The analysis is based on two-way ANOVA with multiple comparison tests (p < 0.01). (**B**) The plaque morphology of rLS1, rLS1-HN-RBD, and rLS1-S1-F was similar. The genetic stability of the rLS1-HN-RBD, and rLS1-S1-F viruses was evaluated at the 3rd and 6th passages by (**C**) western blot analysis and (**D**) RT-PCR. P3: represents the 3^rd^ passage and P6: the 6^th^ passage. The black arrow indicates the expected molecular weight.

The genetic stability of the recombinants virus on 9 days old SPF embryonated chicken eggs sequentially was estimated as 6 passages. Viral genome analyzed by RT-PCR at 3rd and 6th passage did not show any evidence of variability in the size of the gene inserts. Western blot analysis confirmed that the expression of the S1-F and HN-RBD proteins was not affected during passages (Figure 3C-3D).

The IPIC in one-day-old chickens showed that rLS1-HN-RBD (0.0) and rLS1-S1-F (0.0) viruses were slightly attenuated compared to the rLS1 (0.1). The MDTs were 120 h and 118 h for rLS1-S1-F and rLS1-HN-RBD viruses respectively. The MDT for rLS1 was 108 h.

#### 3.1.4. Stability of the lyophilized vaccine

The lyophilized vaccine could be stored at 4°C, and the titers by plaque assay at day 1, 30, and 50 were 8.37± 0.06, 7.69 ± 0.24, and 8.24 ± 0.05 log_10_ PFU/mL respectively. Similarly, the HA titers were identical (2^10^) after the three periods of storage. The expression of S1-F and HN-RBD proteins in Vero E6 cells infected with the lyophilized NDV vaccine was confirmed at day 1, 30, and 50 days post-lyophilized by Western blot assay (Figure 4).

**Figure 4.**
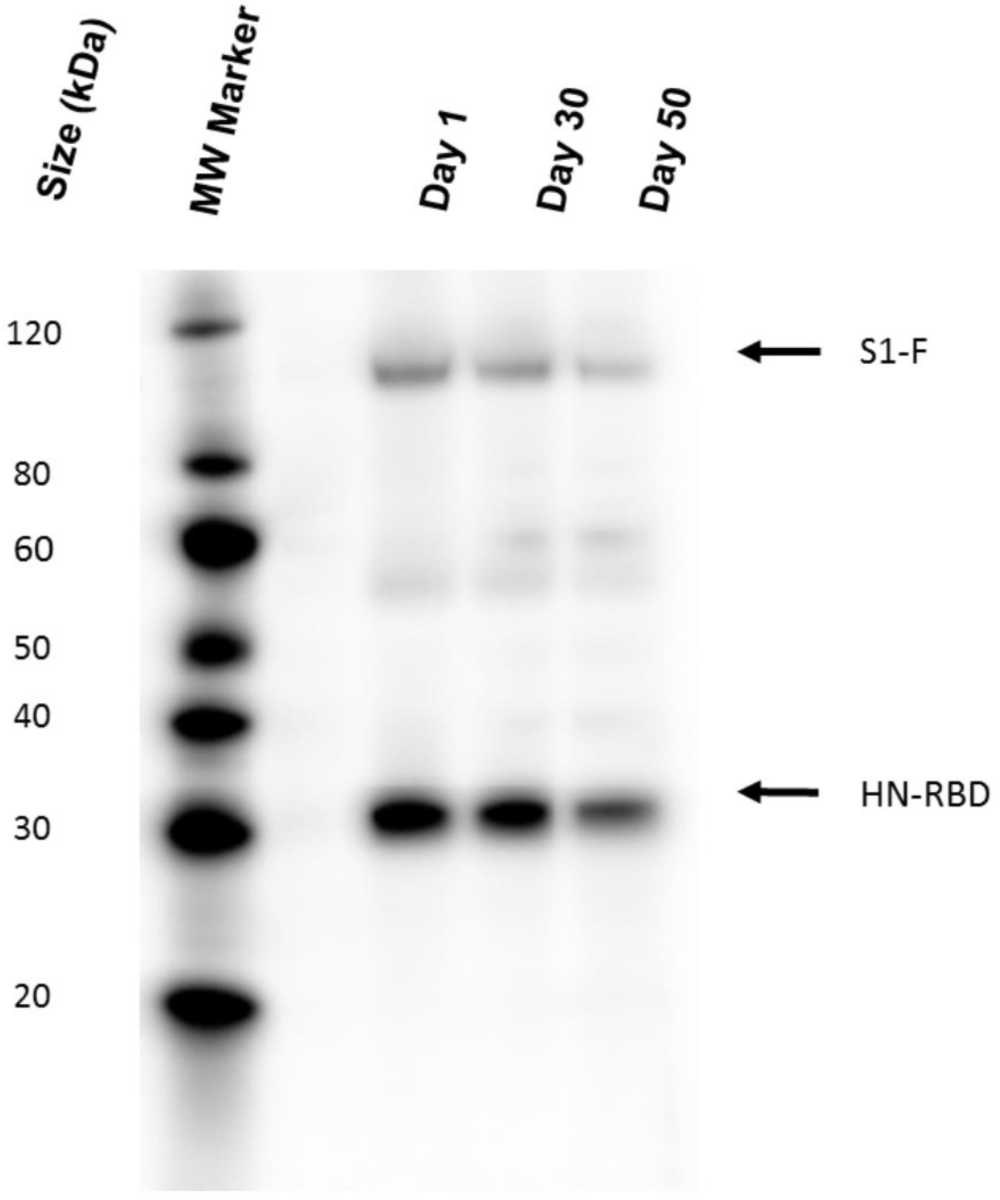
**S**tability of the lyophilized NDV vaccine. The expression of S1-F and HN-RBD proteins in Vero E6 cells infected with the lyophilized NDV vaccine was confirmed at day 1, 30, and 50 days post-lyophilized by Western blot assay using a rabbit antibody specific to SARS-CoV-2 RBD protein and Anti Rabbit IgG conjugated to HRP. The black arrow indicates the expected protein band.

### 3.2. Safety-like assay in mice

Histopathological analyses in the lungs of the 4 mice groups (including the negative control group) inoculated with the three NDV vaccines show no evident signs of lesions in bronchioles, blood vessels or alveolar parenchyma (Figure 5). Although the interstitial spaces in all experimental groups were slightly thickened, this was also observed in the control group. However, slightly celled parenchyma was observed in the groups inoculated with the NDV vaccine compared to that observed in the control group.

**Figure 5.**
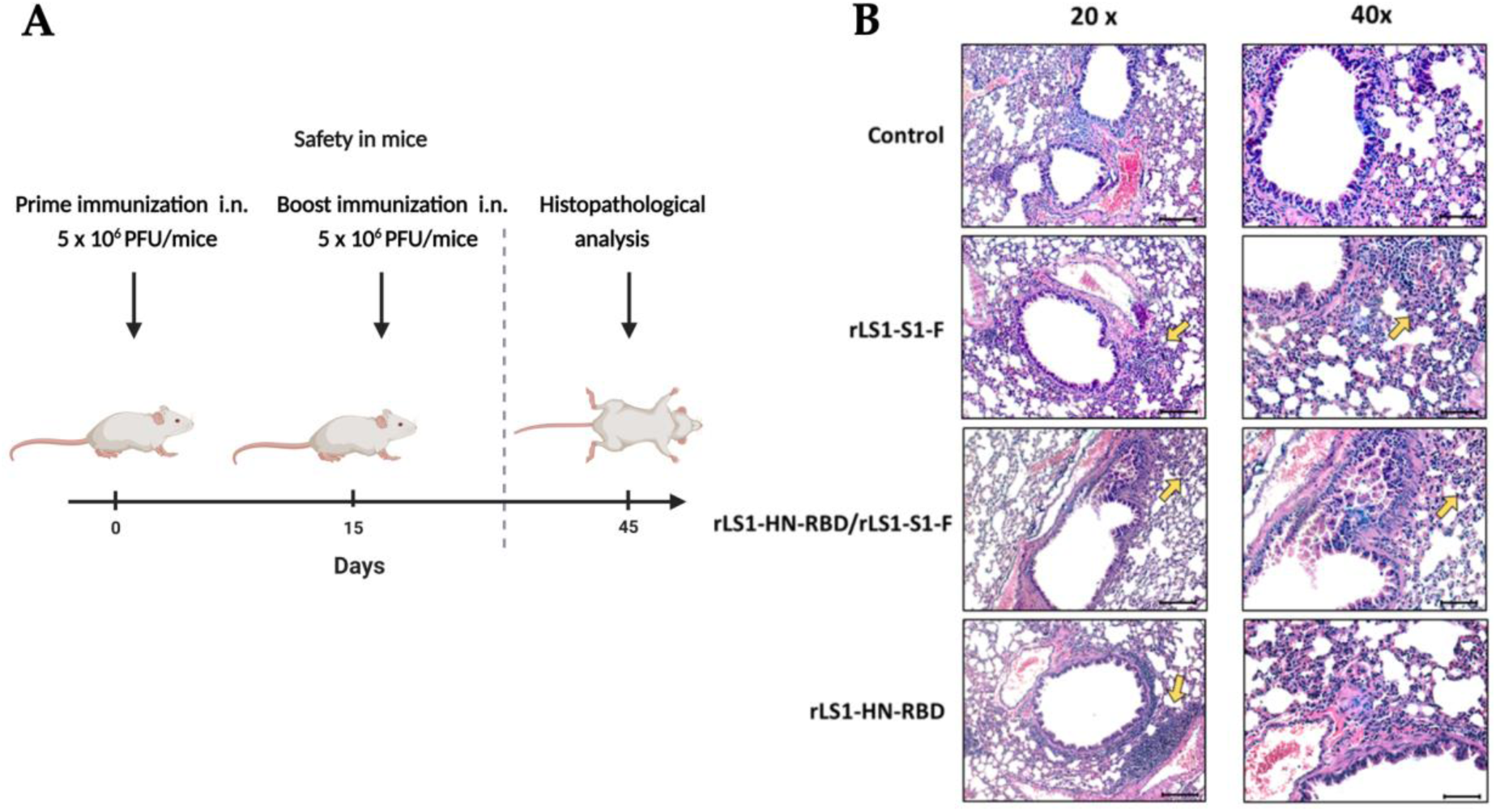
Histopathological lung analysis of mice inoculated with NDV vaccines. (**A**) Inoculation regimen. Mice were immunized by the intranasal route using a prime-boost regimen with a two-week interval. Four groups of mice were included in this study: group 1 (rLS1-S1-F; *n*=4), group 2(rLS1-HN-RBD; *n*=2), group 3(rLS1-S1-F/rLS1-HN-RBD; *n*=4), group 4 (negative control, *n*=4). (**B**) Organs were obtained 30 days after boost and stained with hematoxylin-eosin. Slightly cellular regions are represented by a yellow arrow. Scale bar in 20x amplitude: 100 microns; scale bar in 40x amplitude: 50 microns.

### 3.3. Toxicity-like assay in rats

During the 21 days of the study, the animals received three inoculations intranasally of rLS1-S1-F virus, and no mortality was recorded. The weight variation of the treatment groups and the control group was not significantly different (Figure 6B). No mortality was recorded at 72 hours in any of the treatment groups. The median lethal dose (LD50) value by intranasal route is greater than 1.39×10^9^ PFU/kg.

**Figure 6.**
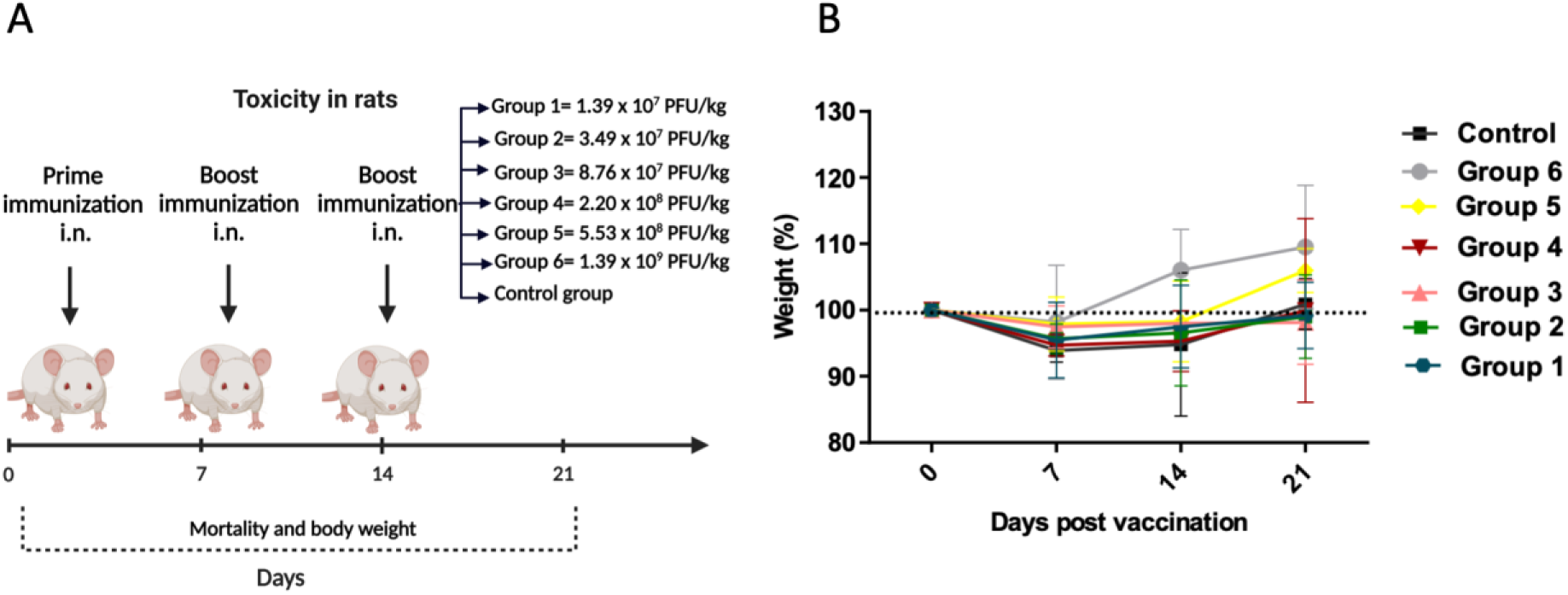
In vivo toxicity in rats. (**A**) Immunization regimen. Male albino rats of the Wistar strain were intranasally immunized with rLS1-S1-F vaccine at days 0, 7, and 14. Seven groups of rats (*n* = 6) were included in this study. (**B**) Rats Bodyweight measure were registered daily during 21 days after intranasal administration of rLS1-S1-F virus at different concentrations: Group 1 (1.39×107 PFU/kg), group 2 (3.49×107 PFU/kg), group 3 (8.76×107 PFU/kg), group 4 (2.20×108 PFU/kg), group 5 (5.53×108 PFU/kg), group 6 (1.39×109 PFU/kg) and the control group (allantoic fluid).

### 3.4. Immunogenicity in hamsters

#### 3.4.1. The intranasal vaccine elicits specific antibodies against S protein and neutralizing antibodies against SARS-CoV-2 in hamsters

Fifteen days after prime immunization, the hamster groups immunized with the live rLS1-HN-RBD, rLS1-S1-F, and the mixture rLS1-HN-RBD/rLS1-S1-F vaccines developed specific serum IgG antibodies against S1 and RBD proteins of SARS-CoV-2. At 15 days post-boost (30 days post-immunization) there was significant increase in the titers of serum IgG antibodies. The control group did not induce SARS-CoV-2 S1 and RBD-specific serum IgG antibody. Immunization with rLS1-S1-F induced a significative higher level of S1 and RBD-specific serum IgG antibody at 15 days post-boost when compared to rLS1-HN-RBD and the mixture of rLS1-HN-RBD/rLS1-S1-F vaccines (Figure 7B-7C).

**Figure 7.**
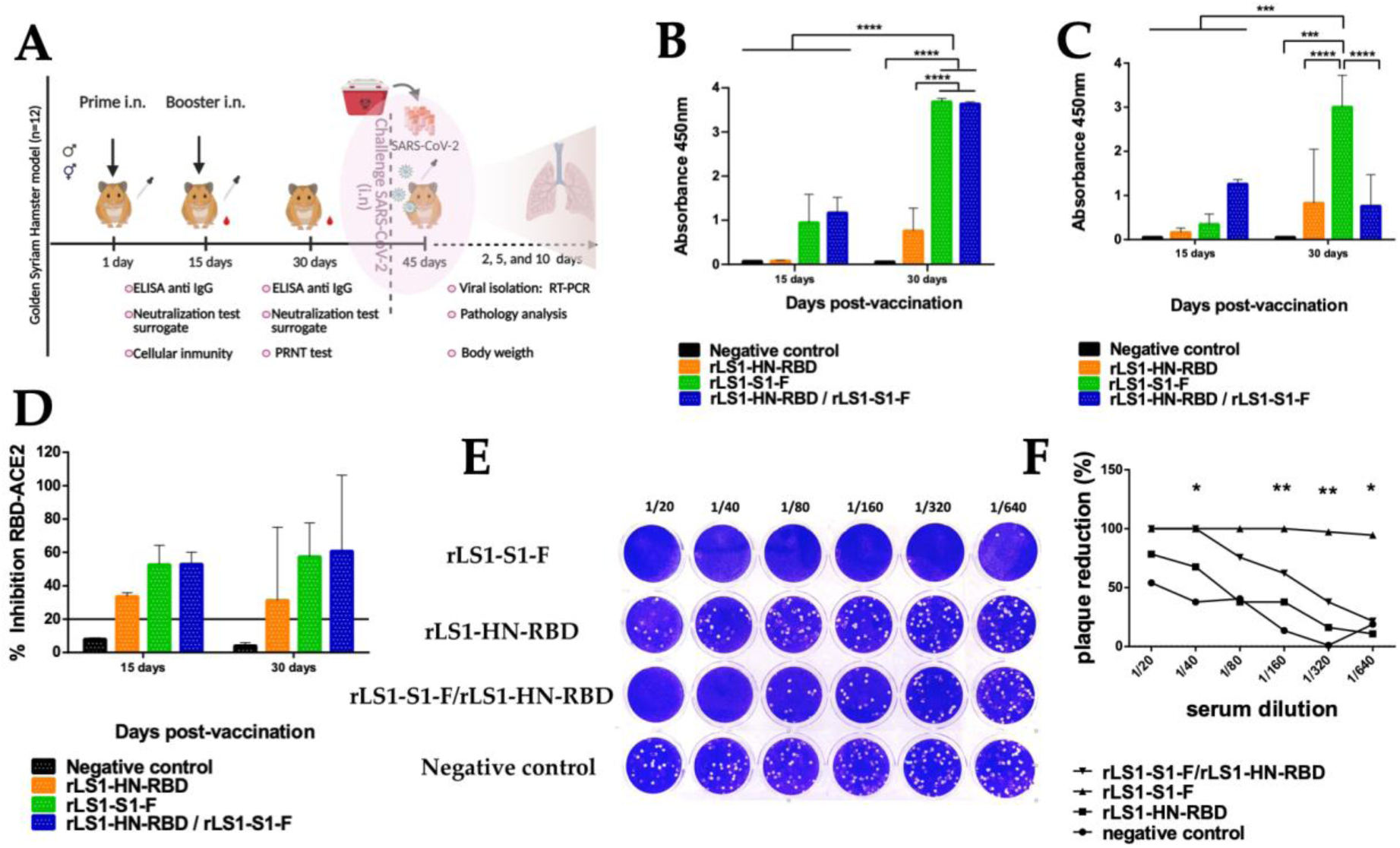
The intranasal vaccine elicits specific antibodies against RBD protein and neutralizing antibodies against SARS-CoV-2 in hamsters. (**A**) Immunization regimen. To evaluate the immunogenicity of the NDV vaccines, five-week-old female and males golden Syrian hamsters were used in this study. The hamsters were randomly divided into five groups. The Hamsters were vaccinated intranasal route with live NDV vaccine following a prime-boost-regimen with a two-week interval. Group 1 received rLS1-HN-RBD (*n*=12), Group 2 received the rLS1-S1-F (*n*=12), Group 3 received the mixture of rLS1-HN-RBD/rLS1-S1-F (*n*=12), Group 4 did not receive vaccine (*n*=12) since group served controls, and Group 5 receive no vaccine and was not challenged, hence serving as healthy control (*n*=12). One boost immunization with the same concentration of each vaccine was applied in all vaccinated groups at the second week. (**B**) ELISA assay to measure SARS-CoV-2 RBD-specific serum IgG antibody, and (**C**) S1 subunit-specific serum IgG antibody. Sera from hamsters at pre-boost and 15 days after boost were evaluated. SARS-CoV-2 RBD purified recombinant protein was used for ELISA. The cutoff was set at 0.06. (**D**). Immunized hamsters were bled pre-boost and 15 days after boost. All sera were isolated by low-speed centrifugation. Serum samples were processed to evaluate the neutralizing antibody titers against SARS-CoV-2 RBD protein using the surrogate virus neutralization test (sVNT). The positive cut-off and negative cut-off for SARS-CoV-2 neutralizing antibody detection were interpreted as the inhibition rate. The cut-off interpretation of results: result positive ≥20% (neutralizing antibody detected), result negative <20% (neutralizing antibody no detectable). (**E**) Figure depicts titers of plaque reduction neutralization test (PRNT) of SARS-CoV-2 on Vero cells with pool serum from hamsters immunized with rLS1-S1-F, rLS1-HN-RBD, and the mixture of both. (**F**) Plaque reduction (%) curves using pools serum from the different groups of hamsters. Two-way ANOVA and Tukey’s post hoc were performed. *: p < 0.05. **: p < 0.01. ***: p < 0.001. ****: p < 0.0001.

The neutralization assays using the surrogate virus neutralization test (sVNT) determined that the sera of immunized groups with rLS1-HN-RBD, rLS1-S1-F, and rLS1-HN-RBD/rLS1-S1-F vaccines developed neutralizing antibodies specific to RBD protein at 15 days post-immunization and 15 days post-boost, where the sera from hamsters vaccinated with rLS1-S1-F and rLS1-HN-RBD/rLS1-S1-F showed a percentage of inhibition of the RBD-ACE2 union higher than 50%. The LS1-HN-RBD group only showed a 30 % of inhibition up to 15 days post-boost. Sera of the control group remained below 20 % up to 15 days post-boost and did not show neutralizing antibodies against RBD protein (Figure 7D).

Pool serum from hamsters vaccinated with the rLS1-S1-F virus showed a strong titer of reduction of viral plaques in the neutralization assay (PRNT) at 15 days post-boost, retaining 100% of this capacity even at higher dilutions of the serum (1/160). The mixture rLS1-HN-RBD/rLS1-S1-F vaccine showed less titer of reduction of viral plaques (1/40), and rLS1-HN-RBD did not show a titer of reduction of viral plaques (Figure 7E-7F).

#### 3.4.2. Cellular immunity: Cytokines quantification by ELISA

For cytokines IL-4 and IL-10, no detectable values were obtained. For TNFα, values were low and only obtained for the treatments rLS1-S1-F and rLS1-HN-RBD/rLS1-S1-F (Supplementary Figure 1). The values obtained in IL-2 (Figure 8A) for each treatment do not present a significant increase for rLS1-HN-RBD (p = 0.554), rLS1-S1-F (P = 0.0756) and rLS1-HN-RBD/rLS1-S1-F (P = 0.0756). However, one of the individuals analyzed which was vaccinated with rLS1-HN-RBD/rLS1-S1-F, presented high levels of circulating IL-2. Regarding IFNγ, (Figure 8B) the values obtained did not show a significant increase for rLS1-HN-RBD (p = 0.083), rLS1-S1-F (P = 1.00) or rLS1-HN-RBD/rLS1-S1-F (P = 0.564). There is no increase in the levels of Th1 cytokines evaluated, so it is considered that immunization with NDV in hamsters does not induce a significant increase in IL-2 and IFNγ in circulating serum evaluated by quantitative ELISA.

**Figure 8.**
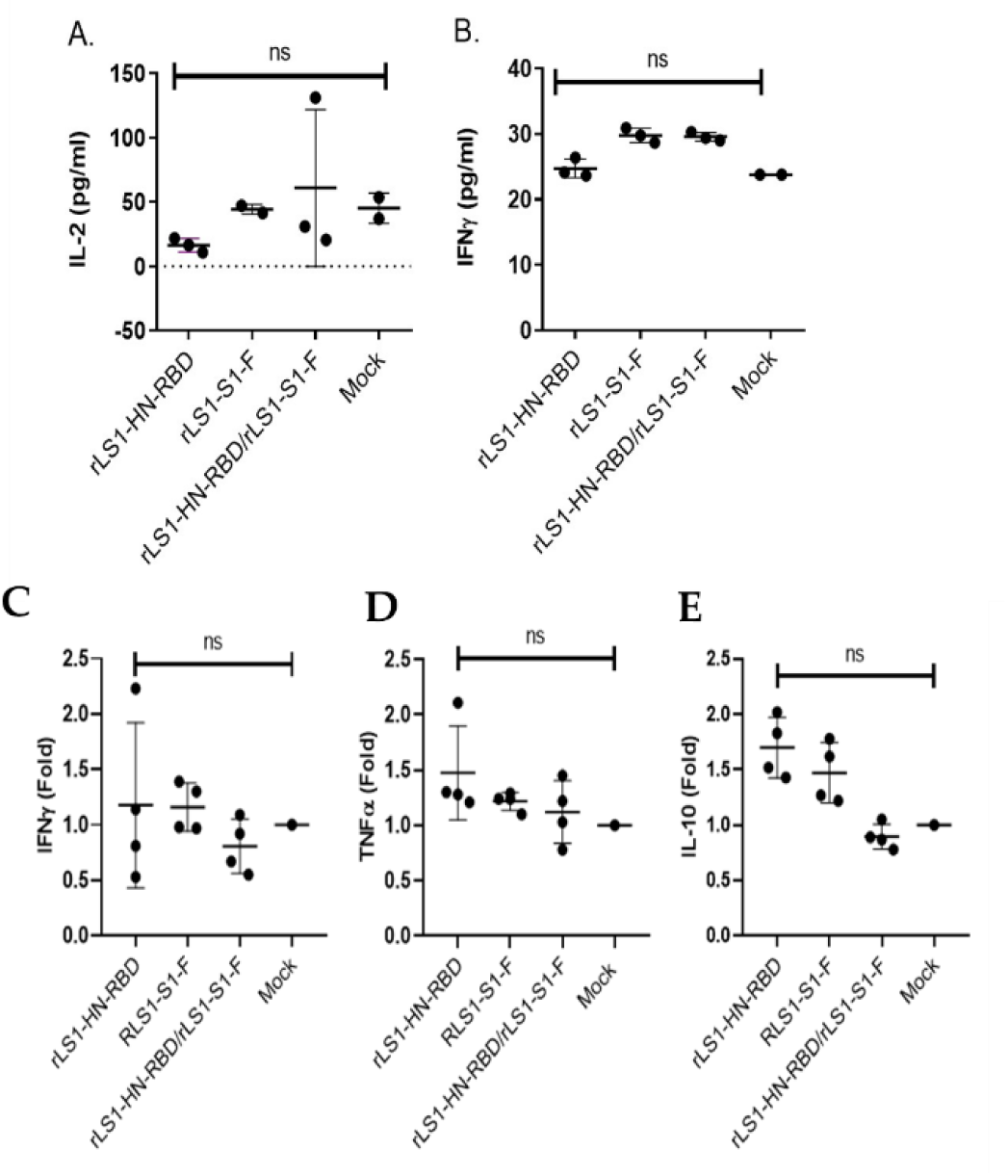
Cellular immunity. These figures show cytokines measured by quantitative ELISA (pg/ml) on hamster serum immunized with rLS1-HN-RBD (n=3), rLS1-S1-F (n=3 for IFNγ, n=2 for IL-2), rLS1-HN-RBD/rLS1-S1-F (n=3) and mock (n=2) at 15 DPV. (**A**) IL-2 and (**B**) IFNγ. ns: not significant; p <0.05. Fold expression of cytokines by ΔΔqPCR from hamster spleens (n=13) vaccinated with rLS1-HN-RBD (n=4), rLS1-S1-F (n=4), rLS1-HN-RBD/rLS1-S1-F (n=4), and mock (n=1). IFNγ (**C**), TNFα (**D**), and IL-10 (**E**), were evaluated at 15 DPV. Each individual present 3 technical replicas for GOI and 2 technical replicas for HKG, a No-RT control was included. Non-parametric Mann-Whitney-Wilcoxon test was used with Stata software v.16. P values of <0.05 were considered significant. * P <0.05, ** P <0.01, *** P <0.001, **** P <0.0001. NS, not significant

#### 3.4.3. Cellular immunity: Cytokines quantification by qPCR

No significant difference in expression was observed for rLS1-HN-RBD (IFNγ 1.2-fold, P=1.000, TNFα 1.48-fold, P=0.1573, IL-10 1.7-fold, P=0.1573), rLS1-S1-F (IFNγ 1.16-fold, P=0.1573, TNFα 1.48-fold, P=0.1468, IL-10 P=0.1573) or rLS1-HN-RBD + rLS1-S1-F (IFNγ 0.81-fold, P=0.4795, TNFα 1.12-fold, P=0.4795, IL-10 0.90, P=0.4795). Regardless of the vaccine, there was no increase in the cytokine levels (Figure 8C-8D-8E).

### 3.5. Efficacy of the vaccines against SARS-CoV-2 challenge

On days 2 and 5 post-challenge with the SARS-CoV-2, the control group not vaccinated, showed a positive isolate (100% of the hamsters) of the SARS-CoV-2 in Vero cells, and it disappeared on day 10 (0% of the hamsters). Viral isolation and IFA showed that rLS1-S1-F and mixture rLS1-S1-F/rLS1-HN-RBD vaccine elicited the best response evidenced by the negative isolation of the virus, and negative IFA detection at days 5 and 10 (0% of the hamsters) post-challenge in the lung tissue. The rLS1-HN-RBD vaccine alone does not have enough neutralizing capacity to stop the infection. It showed a positive isolate confirmed by IFA in lung tissue at day 5 post-challenge (100% of the hamsters) (Figure 9A).

**Figure 9.**
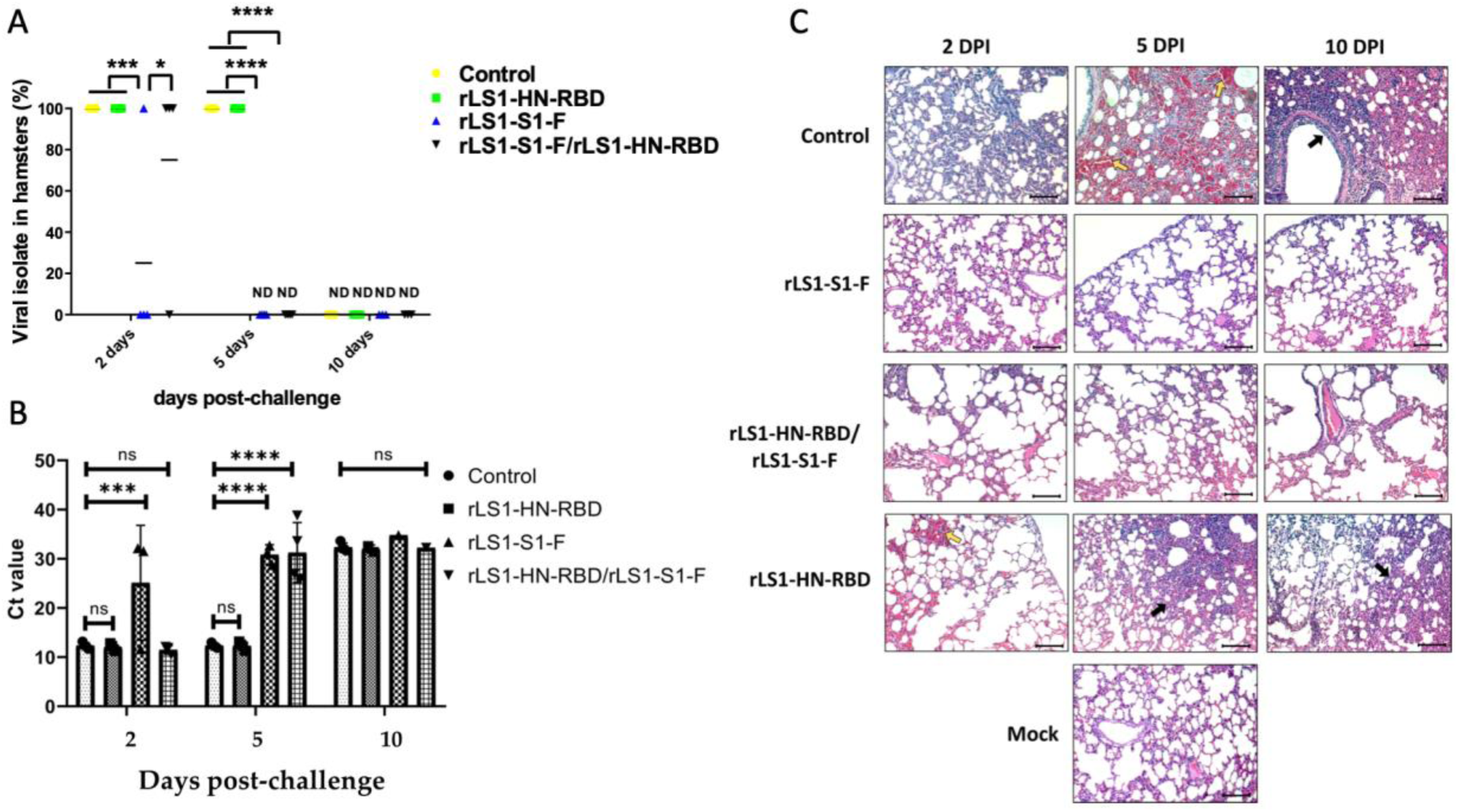
Efficacy of live NDV vaccines against SARS-CoV-2 infection in hamsters. Golden Syrian hamsters groups vaccinated with rLS1-S1-F, rLS1-HN-RBD, the mixture rLS1-S1-F/rLS1-HN-RBD, and negative control (not immunized) were challenged 30 days after the boost with SARS-CoV-2, also a not immunized and not challenge group was included (Mock). (**A**) Viral isolate (%) was done from the lung of each hamster group (n=4) at days 2, 5, and 10 post-challenge. Two-way ANOVA and Tukey’s post hoc were performed. *: p < 0.05. **: p < 0.01. ***: p < 0.001. ****: p < 0.0001. (A) Detection by qRT-PCR of SARS-CoV-2 in culture supernatant of Vero cells, infected with immunized and challenged hamster lung homogenates. The data showed a significant difference in the Ct value * P <0.05, ** P <0.01, *** P <0.001, **** P <0.0001. NS, not significant. (**C**) Lung histopathology of each hamster group (n=4) was euthanized at different days post-infection (DPI). Hemorrhagic and infiltrated areas are indicated by a yellow and black arrow, respectively. Image amplitude: 20x. Scalebar: 100 µm.

The RNA copy number of SARS-CoV-2 by RT-qPCR from the viral isolate in Vero cells of hamster lung homogenates confirmed the presence of a high viral load (Ctp: 12) present in the viral isolate from the lungs of hamsters not immunized (control) and immunized with rLS1-HN-RBD and the mixture rLS1-HN-RBD/rLS1-S1-F vaccines at days 2 post-challenge. Animals immunized with rLS1-S1-F showed a lower viral load (Ctp: 25). At days 5 post-challenge, the viral load from the isolate was maintained in the control group and immunized with rLS1-HN-RBD, however, in hamsters immunized with rLS1-S1-F and rLS1-HN-RBD/rLS1-S1-F the load decreased viral load was significant (Ctp: 28-31) with a p-value <0.05, as shown in Figure. On day 10 post-challenge, all the groups analyzed, including the control, presented a low viral load from viral isolate (Ctp: 32-34) (Figure 9B), probably due to the presence of residual RNA, since some samples, no viral load was detected and we did not detect a cytopathic effect in the viral isolate in Vero cells. The histopathological status of the hamster lungs was monitored during the challenge with SARS-CoV-2. The challenged control group demonstrated the pathological signs of the disease, starting with interstitial pneumonia at 2 days post-challenge, evolving into hemorrhagic pneumonia at 5 days post-challenge, and ending up in severe bronchopneumonia, characterized by a thickening in the parenchyma wall, greater infiltration of inflammatory cells, and in bronchioles lumen, as the loss of alveolar architecture. The groups immunized with rLS1-S1-F and rLS1-HN-RBD/rLS1-S1-F vaccines did not show visible lesions, maintaining the characteristics of the lung tissue as the unchallenged group during all the evaluated times. However, the group immunized with rLS1-HN-RBD vaccine developed pathological signs, although to a lesser degree than the control group presenting pneumonia at 2 days post-challenge and ending in a moderate to severe pneumonia at 10 days (Figure 9C).

Animals belonging to the unchallenged control group showed an average percentage variation in weight of no more than 3% over the 10 days of analysis. On the other hand, the control animals (non-vaccinated) that were infected with the SARS-CoV-2 virus showed an average weight loss below 5% on day 5, and below 10% on day 10, corresponding to a significant difference between the unchallenged control group and the challenged control group at days 5 and 10. Concerning the vaccinated groups, there were no statistically significant differences between them and the unchallenged control groups on days 2 and 5, but there was a significant difference between the challenged control group and the rLS1-HN-RBD/rLS1-S1-F group at day 10 after the trial started (Figure 10A). The average speed, average acceleration, and average displacement confirmed that hamsters vaccinated with the mixture rLS1-S1-F/rLS1-HN-RBD are the most mobile on day 5 (time at which symptoms appear) and day 10 post-challenge (time at which symptoms disappear). Considering the mobility parameter, the mixture rLS1-S1-F/rLS1-HN-RBD vaccine allowed a better recovery (Figure 10B-10C-10D).

**Figure 10.**
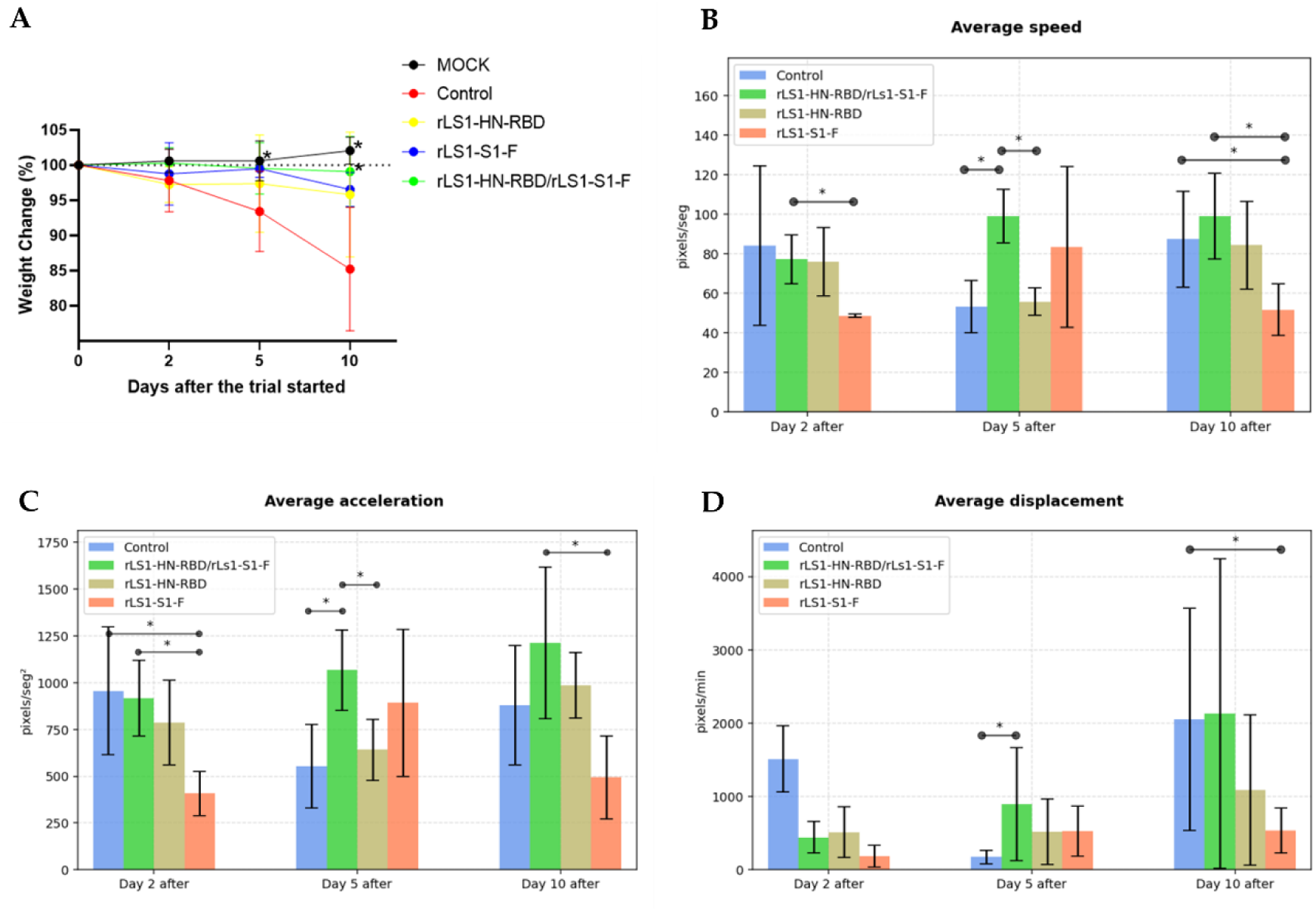
Body weight and mobility analysis of SARS-CoV-2 challenged golden Syrian hamsters. (**A**) Changes in body weight (percent weight change compared to day 0) of hamsters inoculated with SARS-CoV-2 and Mock group, at days 2, 5, and 10 post-challenged. Mobility assessment results shown (**B**) average velocity, (**C**) average acceleration, and (**D**) average displacement. Mean ± s.d. are shown. Asterisks indicate that results were statistically significant compared to the control group (P<0.05).

## 4. DISCUSSION

In this study, we developed two recombinant NDV nasal vaccine candidates expressing the SARS-CoV-2, S1 and RBD antigens. The vaccine candidate expressing S1 showed favorable results during its evaluation in the pre-clinical phase. It showed to be safe according to histopathological analysis in different organs in mice, as well as to the toxicity assay in rats. The efficacy assay in hamsters showed evidence that the vaccine was able to protect the animals against the SARS-CoV-2 virus. The lungs did not show evidence of cell damage, and the viral load as well as the viral viability was considerably lower in the vaccinated group compared to the control group. Confirming these results, the vaccinated animals showed significantly greater mobility than the control group, and showed no evidence of body weight loss, in contrast to what was observed in the control group. Together with these encouraging results, the fact of being a nasal vaccine easy to administer, being lyophilized and stable on storage at 4°C, and being a relatively fast and economical vaccine to produce, make this vaccine candidate a promising tool to contribute in the fight against the COVID-19 pandemic. The two recombinant NDV, the rLS1-HN-RBD and the rLS1-S1-F, showed biological properties, pathogenicity, and growth dynamics, similar to the parental rLS1. Evidence supports that the RBD and S1 subunits were incorporated into NDV particles, showing viral surface expression when fusing these genes with TM/CT from HN and F of NDV, respectively ^49, 50, 68^. This was confirmed by Western blot and flow cytometry analysis. Furthermore, a high-level of expression of RBD and S1 subunits in the infected cells/lysates suggests that both genes were inserted between P and M genes, an optimal insertion site in NDV ^69^, in the 3’-proximal locus of NDV ^70^.

The RBD domain, and in particular the RBM motif, are the regions of the spike protein, which directly interact with the ACE2 receptor to initiate the infection process, thus representing the most important target for neutralizing antibodies ^71, 72^. Recent studies have shown that in COVID-19 convalescent patients, neutralizing antibodies are commonly directed against specific epitopes of the RBD domain^71, 72^.

Three nasal vaccine candidates were evaluated in this study. One was an NDV presenting the RBD domain (rLS1-HN-RBD), the second was an NDV presenting the subunit S1 (rLS1-S1-F), which comprises the RBD domain, and the third was a mixture of both (rLS1-HN-RBD and rLS1-S1-F). The strongest immunity and protection were elicited by the rLS1-S1-F vaccine, followed by the mixture rLS1-HN-RBD/ rLS1-S1-F vaccine. Surprisingly, the rLS1-HN-RBD vaccine did not show evidence of protection. There are at least two possible non-excluding explanations. First, the S1 subunit may include protective epitopes in addition to those present in the RBD domain, for which neutralizing antibodies could interfere with the S-ACE2 interaction perhaps at a distant steric level.

This possibility is supported by a previous study, which showed evidence that the subunit S1 contains neutralizing epitopes not found in the RBD ^73^. Second, that the RBD present in the rLS1-HN-RBD virus did not reach a folding close enough to the 3-dimensional structure of the biologically active RBD when it is in the SARS-CoV-2 viral particle. It is likely that the presence of the additional S1 moiety may provide support for RBD to reach a proper folding. Therefore, it is possible that conformational B epitopes may be playing a more important role than linear T epitopes in the protective immune response. A recent study that evaluated a similar vaccine candidate vectored in NDV exposing the complete S antigen (S1 and S2 domains), confirmed protection of hamsters in a challenge assay ^54^. It is likely that the major contribution to the protection observed in these studies is associated to S1, which includes the RBD domain in a protein environment that favors the proper folding, allowing the presentation of appropriate conformational B epitopes. Animals from the control non-vaccinated group challenged with SARS-CoV-2 showed a positive virus culture at 2 and 5 days post infection (dpi), and it became negative at day 10. This is consistent with previous studies, which reported that viral load is reduced to undetectable levels by 8 days after infection in the hamster animal model ^74, 75^. Along with the culture results, the IFA test also confirmed that the rLS1-S1-F vaccine followed by the mixture rLS1-HN-RBD/ rLS1-S1-F, induced the strongest protective responses, evidenced by lack of virus isolation in the lungs at 5 dpi. At 2 dpi, only half of the animals vaccinated with S1 had virus isolation and negative IFA. This suggests that the inactivation of the inoculated SARS-CoV-2 virus would be occurring between 2 and 3 dpi in the hamster model.

The histopathological evaluation of the non-vaccinated hamsteŕs lungs after the challenge, revealed serious pathological signs of the disease, beginning with interstitial pneumonia 2 days dpi, evolving to hemorrhagic pneumonia and ending in severe bronchopneumonia, as well as the loss of alveolar architecture. In contrast, the groups immunized with rLS1-S1-F and the mixture rLS1-HN-RBD/rLS1-S1-F seemed protected, showing almost intact alveoli, capillaries, and respiratory capillaries, without evidence of an inflammatory reaction. However, the animals vaccinated with rLS1-RBD-HN developed pathological lesions, although with a lesser hemorrhagic degree than the non-vaccinated control group, with signs of pneumonia at 2 dpi and ending in a moderate to severe pneumonia at 10 dpi.

Recent studies reported that Golden Syrian hamsters inoculated by the intranasal route with 8×10^4^ TCID_50_ with SARS-CoV-2 show at 2 dpi, efficient virus replication in the respiratory epithelium associated with a reduction in the number of replicating olfactory sensory neurons of the nasal mucosa^75^. However, virus clearance is observed from 7-10 dpi ^74, 75^. The ability of the rLS1-S1-F vaccine candidate to neutralize the SARS-CoV-2 virus, block its replication in the cell culture between 2 and 5 dpi, and reduce its presence in the lungs according to the IFA assay, suggests the possibility that the rLS1-S1-F vaccine candidate may eventually reduce the virus transmission since 2 dpi in the hamster model. This is an encouraging result, that needs to be evaluated in human clinical trials. In the hamster challenge trial, the animals of the non-vaccinated control group, lost significant body weight (between 10-25%) during the 10-day trial period. Animals vaccinated with rLS1-S1-F showed a body weight variation similar to the corresponding pattern of the uninfected MOCK group, suggesting a protective effect of this vaccine candidate.

In this study, we evaluated for the first time, the mobility pattern of the animals as an objective and quantitative indicator of their health status. From recorded videos, using computational tools of pattern analysis and digital tracking, we measured the average velocity, the average acceleration, and the average displacement of the animals in their cages at 2, 5, and 10 dpi. The results showed that at 5 dpi, the animals from the non-vaccinated control group have a lesser average displacement, velocity and acceleration, compared to the vaccinated animals. This marked difference is not clearly observed at 2 dpi, supporting that at this moment, the infected animals are close to an asymptomatic stage. Similarly, at 10 dpi all animals recovered their mobility with no noticeable differences. This result agrees with previous findings that show SARS-CoV-2 clearance at 10 dpi.

A favorable result that suggests a strong protection of the rLS1-S1-F vaccine candidate, is the neutralization capacity of the vaccinated hamster’s sera against the SARS-CoV-2 virus at 30 days after immunization, in the PRNT test. In contrast, animals vaccinated with rLS1-HN-RBD showed marginal sero neutralization capacity. This result agrees with similar findings reported in a recent study evaluating an intranasal NDV live attenuated vaccine, showing that humoral immunity is induced to high levels in a relatively short time ^76^.

Respiratory viruses induce a strong response in the respiratory tract mucosa, so vaccines based on the NDV vector should be effective in reducing transmission of respiratory diseases. Likewise, a vaccine candidate for SARS-CoV based on the NDV construct with the S protein confirmed that two doses every 28 days of 10^7^ PFU by the intranasal and intratracheal routes, produced significant protection against SARS-CoV in juvenile primate models, *Cercopithecus aethiops* ^42^. In a recent study, it was shown that an NDV-vectored nasal vaccine, displaying the S protein, was able to induce an IgA antibody-mediated mucosal humoral response ^76^. A mucosal antibody response is considered important against infections that use the respiratory tract as a route of entry, making the respiratory tract mucosa, the first line of defense against this infection. At the moment of presenting this manuscript, we are completing an assay to evaluate the presence of anti-S1/RBD mucosal IgA antibodies in mice vaccinated with rLS1-S1-F. A preliminary result indicates evidence of IgA production in bone marrow cells (data not shown). Further studies are required to confirm this finding. An optimal immunogenic response in vaccinated animals should include T cell immunity ^77^ with Th1 response, which is in particular necessary for a successful control of SARS-CoV-2 ^5, 76^. Previous studies found that Th1 response was found as a systemic T-specific IFNγ response after intranasal vaccination in mice at 19 days post immunization ^5, 76^. To evaluate the cellular immune response induced by our NDV-vectored vaccines, we measured the expression levels of cytokines IFNγ, TNF-α and IL-10 in the spleen of vaccinated hamsters using qPCR. This approach was previously used for cytokine evaluations after challenge with SARS-CoV-2 ^61, 62, 78, 79^. In our study, a trend for an increase of cytokines production was observed for IL-10 and TNF-α in vaccinated hamsters, although it was not statistically significant. It is likely that this lack of significance could be attributed to an insufficient sample size or to the fact of having analyzed specimens of 15 days after immunization. Further studies are needed to confirm this observation. The detection of Th1 cytokines (IL-2 and IFNy) in circulating serum by ELISA was limited by a reported low sensitivity, and by the lack of reagents to evaluate immunogenicity in Golden Syrian hamsters ^78^. A recent study suggested that a vaccine that stimulates IL-2 with low expression of IL-10, is likely to be activating the Th1 response_80._

The safety of the three vaccine candidates was evaluated in a murine model, through histopathological evaluation of the lungs and other organs after vaccination. Our results showed that mice immunized with the NDV-vectored vaccine candidates did not show evidence of damage in the different organs evaluated by histopathological analysis. No lesions or obstructions were observed, neither in the lumen of the bronchioles nor in the blood vessels. Although slightly cellular parenchyma is present, this could correspond to the stimulation of T cells, which is a temporal phenomenon already described for this type of vaccines ^76^. In addition, the evaluation of toxicity demonstrated that the rLS1-S1-F vaccine candidate can be delivered in up to 3 doses (one dose more than planned), even at a concentration 100-fold higher than planned. Likewise, the slightly decrease of body weight in both the challenged and control groups, is within the limits of weight loss previously reported in similar studies ^81, 82^. Nevertheless, the weight loss could be attributed to the possible stress of administering the dose, and/or the need of more acclimatization days. However, it is important to remark that one week after the booster, an increase of body weight was observed in all groups. Likewise, no visible lesions were observed in the nasal area, the site of administration of the vaccine candidate, nor any notable clinical symptom or evidence of mortality was recorded. These results are comparable with those obtained in the different pre-clinical studies of currently available SARS-CoV-2 vaccines ^83^.

Other studies have reported that NDV virus is restricted to the respiratory tract, and is not detected in other organs or blood. Therefore, replication in humans is expected to be limited and benign ^84, 85^. Rarely, humans exposed to mesogenic NDV have been observed to develop conjunctivitis, laryngitis, or flu-like symptoms that disappear within 1 to 2 days ^85, 86^. It is important to remark that the safety and toxicity tests were conducted with doses that were not produced under strict GMP certification. Having conducted these trials with non-GMP doses of the vaccine candidates resulted in evaluating them in a more adverse scenario than expected. A favorable outcome in such an adverse scenario strongly suggest that it would be replicated when testing the vaccine doses produced under GMP conditions.

NDV has been reported over the years as a tool for successful vaccine development to eliminate infectious diseases in poultry ^50^. Currently, there is a NDV strain genotype XII that predominates in Peru, China and Vietnam, which has been successfully used to develop rNDV vaccines ^87^. Likewise, Shirvani et al. used the rNDV vector to control infectious bronchitis in free-raising chickens, in which the spike S protein is expressed, showing equally efficacy to the same infection bronchitis virus vaccine ^49^.

LaSota is the most common attenuated NDV strain used as a vaccine around the world against NDV infection, with doses of TCID_50_ ranging from 10^4^ to 10^5^, that is administered to animals by oral, nasal, ocular or by spraying delivery ^88–90^. This results in frequent exposure of vaccinators to the NDV virus, which ends up being inhaled. Thus, vaccinators and caretakers are routinely exposed to the NDV vector, without presenting any side effects to date ^39, 91, 92^. There have not been reports of adverse effects by the accumulated doses of NDV inhaled by vaccinators in Chincha-Peru over the last decades (data not shown).

For the past 10 years, NDV has been used as a recombinant oncolytic and immunotherapeutic agent in humans, administered intravenously, inducing long-lasting immunity in different types of cancer. The resistance of healthy cells to NDV infection is explained by the overexpression of the antiviral genes RIG-I, IRF-3, IFN-β, and IRF7. In contrast, tumoral cells have a low expression of these genes, which make them susceptible to NDV. In tumoral cells, NDV replicates in the intra cellular environment with a cytotoxic, necrotic effect or an auto phagocytosis mechanism. Thus, the NVD achieves control in cancer cells, stimulating immunity against tumors ^45, 93^. It has been shown that NDV is non-pathogenic to humans as it has been used as a treatment against multiple cancer types without causing serious adverse effects ^45, 93^. In vitro studies in breast, kidney, bone, skin, brain and cervical cancer cells show that NDV causes cancer cell reduction by activating extrinsic and intrinsic apoptosis pathways ^45, 94–97^. Clinical trials with rNDV have been developed over the years, and more than a dozen of studies have been reported in which lytic NDV strains have been inoculated intravenously, intratumorally or intramuscularly. For example, a phase III against colon cancer has shown an increase in the lifespan of volunteers in 7 years, without severe side effects ^48^. Likewise, phase I has been carried out in patients with chronic cancer, using the mesogenic strain rNDV PV701, resulting in the survival of volunteers with up to two more years of life ^45^. The rNDV has also been tested in phase I and II against malignant gliomas, where the doses inoculated intravenously for 5 consecutive days ranged at concentrations of 0.1, 0.32, 0.93, 5.9, and 11 BIU of NDV-HUJ in which patients presented a mid-fever as a side effect that was controlled after a few days ^43, 45^. Similarly, clinical studies of NDV in metastatic cancer patients show important evidence of its effectiveness ^43, 98–101^. Because of this, NDV has been proposed as a potential treatment against numerous types of cancer. Additionally, the phase II study in multiform gliomas showed that four of the patients, treated with intravenous NDV, survived up to 9 years ^99^. Likewise, the use of NDV with Durvalubmab (Medimmune/AstraZeneca) has been proposed against various advanced neoplasms (NCT03889275)^99^. In another clinical study, inhalation of approximately 1×10^8^ PFU of NDV was associated with only one day of low fever in 25% in non-human primates ^84^. In addition, it is noteworthy that the NDV-vectored vaccine presented here does not require adjuvants to booster the immune response. It has been shown that the use of nasal adjuvants could result in a lower immune response ^102^, as well as trigger adverse effects, making them unsafe for human use ^103–106^.

The present NDV vectored vaccine candidate has been developed including a final lyophilization step. This confers an important stability, being able to be stored at 8°C for several months without losing more than 5% of its activity, as seen in other lyophilized vaccines ^107^. It is important to highlight that in circumstances where a batch of vaccine is to be used quickly, it could be distributed in liquid medium, reducing the cost significantly. We have observed that the NDV vaccine candidate viruses rLS1-HN-RBD and rLS1-S1-F can be stored in liquid solution for 15 days without losing more than 5% of their activity, measured in PFU units (data not shown).

The rLS1-S1-F vaccine candidate described in this work has been produced in SPF embryonated chicken eggs. However, it is also possible to produce it with the same or better efficiency in cell culture using single-use bag-based bioreactors. We have recently standardized a protocol that allows us to produce the rLS1-S1-F vaccine candidate in a bioreactor, achieving titers similar to those achieved in SPF embryonated chicken eggs (data not shown). A single 1000 Lt bioreactor based a single-use system could produce 1 million doses approximately every 72 hours. This means that the production of the rLS1-S1-F vaccine candidate could be even more efficient, with a cost per dose of a fraction of a dollar (data not shown).

The fact that the NDV-S1 vaccine is administered through the nasal route gives it a notable advantage in simplifying the logistical requirements for immunizations. There is no need for an army of vaccinators or large numbers of syringes. It is possible that doses of rLS1-S1-F vaccine could be delivered in 500-dose vials with a manual trigger-activated dispenser system that uses individual disposable tips. In this way, a nasal vaccine could be delivered in large-scale campaigns in remote rural communities with great ease.

The COVID-19 pandemic may become endemic. So, if it were to happen, vaccines would need to be routinely administered with some frequency ^108,109^. SARS-CoV-2 in recent months has shown an intense level of mutations in the viral antigens used in the various vaccines currently available. These mutations have been selected naturally, in the face of the immunological pressure exerted by individuals cured of COVID-19. Thus, mutations have now been identified to give the virus the ability to escape acquired immunity (immune resistance), and thus the increased frequency of COVID-19 reinfections. These same naturally selected mutations have also been selected in-vitro, under immunological selection pressures using convalescent sera neutralizing antibodies. This suggests the possibility that vaccines based on the S1 antigen circulating in the early 2020’s could have their effectiveness compromised against the new SARS-CoV-2 variants. Therefore, an efficient way to deal with the COVID-19 pandemic will be to use vaccines customized for specific geographic areas that can be produced and administered promptly, based on the distribution of circulating variants over a certain period of time. The rLS1-S1-F vaccine candidate can be updated to carry a vaccine antigen corresponding to a new relevant strain in a relatively short time. In 30-45 days, a NDV could be transformed and a master cryobank generated to start producing updated vaccine batches. It is therefore important to have permanent epidemiological surveillance programs to identify any variation in the distribution of circulating strains.

*In conclusion*, we demonstrated that rLS1-S1-F vaccine candidate showed positive results in the pre-clinical studies. This vaccine candidate showed evidence to be safe and immunogenic, and provides strong protection against SARS-CoV-2 in the infection challenge. Clinical trials should be conducted to evaluate its safety and efficacy in humans.

## ACKNOWLEDGEMENTS

We thank the National Institute of Health from Peru (INS) for providing the SARS-CoV-2 virus aliquots necessary for the challenge assay, as well as for their participation in the virus neutralization, viral load and viability tests. In particular, we acknowledge Lic. TM Maribel Carmen Huaringa Nuñez and BSc. Pamela Lisset Ríos Monteza.

We acknowledge Dr. Maria Salas, for her advice in the toxicity study.

We are grateful to Dr. Valerie Paz-Soldan, Dr. Gabriela Salmon, Dr. Danni Kirwan, and Dr(c) David Requena for their revisions and criticisms to the manuscript.

## PATENT

The vaccine candidates presented in this study have been filed for patent request. Patent pending N33-2021/DIN.

## Supplementary Material

**Supplementary Table 1.**
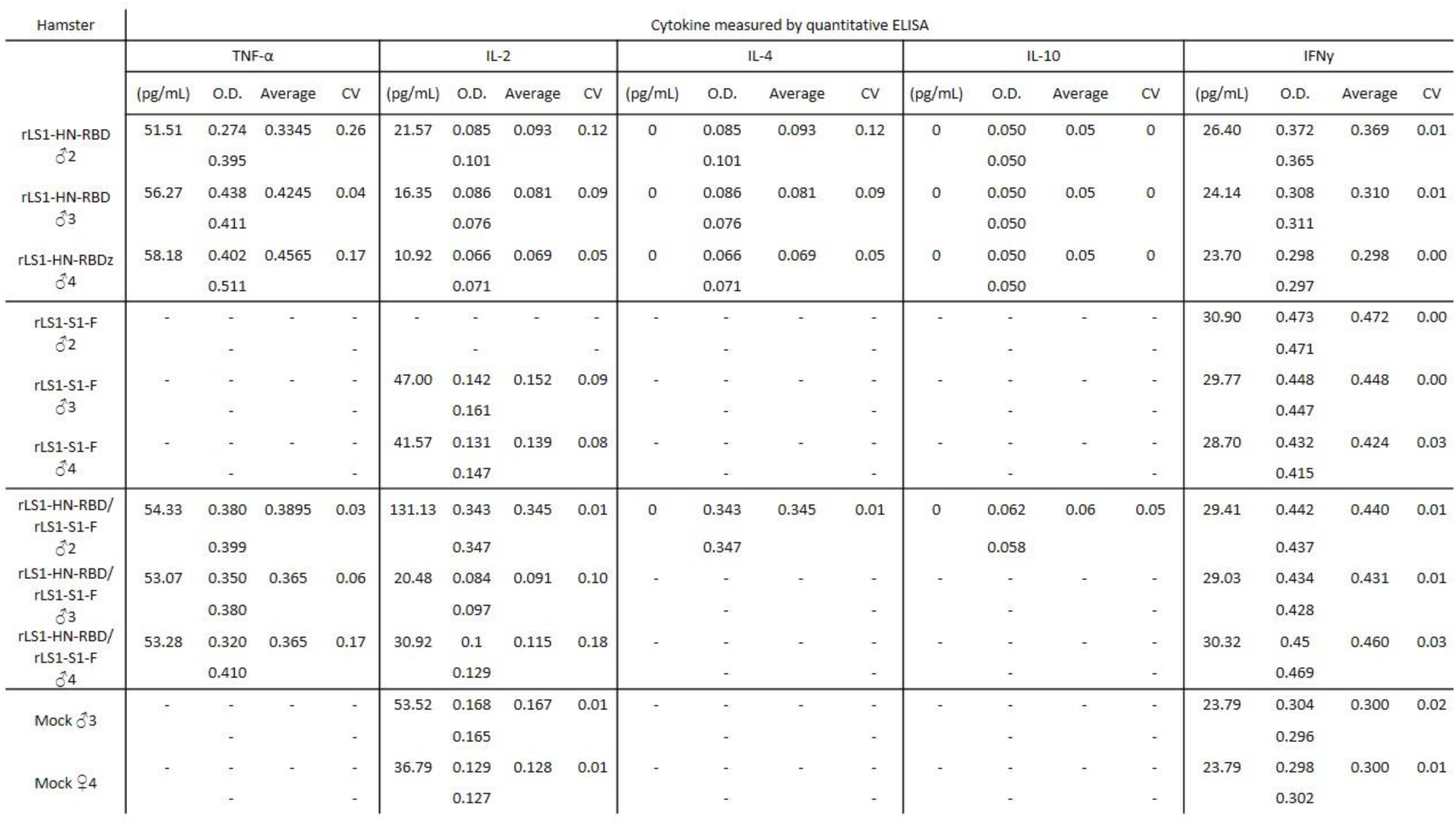
Concentration of cytokines (pg/mL) of hamster serum 15 DPV with rLS1-HN-RBD (n=3), rLS1-S1-F (n=3 for IFNγ, n=2 for IL-2), rLS1-HN-RBD/rLS1-S1-F (n=3) and mock (n=2). The sera were placed in duplicate in 96-well plates precoated with antibodies against the hamsteŕs cytokines, incubated at 37 °C and revealed with the enzyme streptavidin or avidin-HRP, to finally add the substrate TMB, stopping the reaction with sulfuric acid. The plates were developed in the spectrophotometer at 450 nm.

**Supplementary Figure 1.**
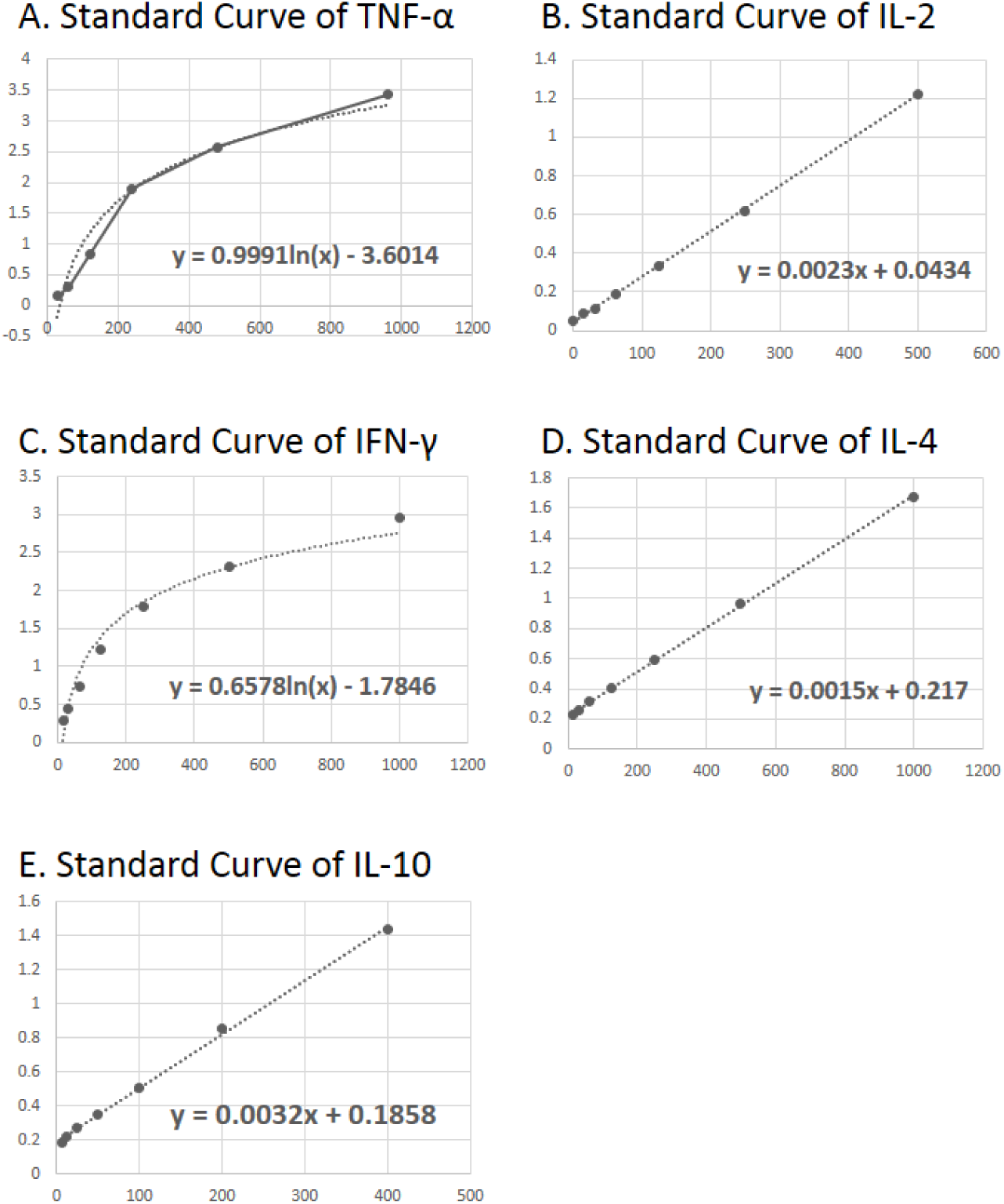
Standard Curves. The standard curve was constructed following the manufacture’s indications (MyBiosource.com). The plaque was precoated with the antibodies and was incubated with the serial dilution of standard or with the standard included in the kit, for 2 h at 37°C, followed by extensive washes. The plaque was developed using streptavidin-HRP and TMB solution, and sulfuric acid was used as stop solution. The plaque was read in the spectrophotometer at 405 nm, and the standard curve was made using the theoretical concentration of each standard established in the manufacturer instruction.

**Supplementary Figure 2.**
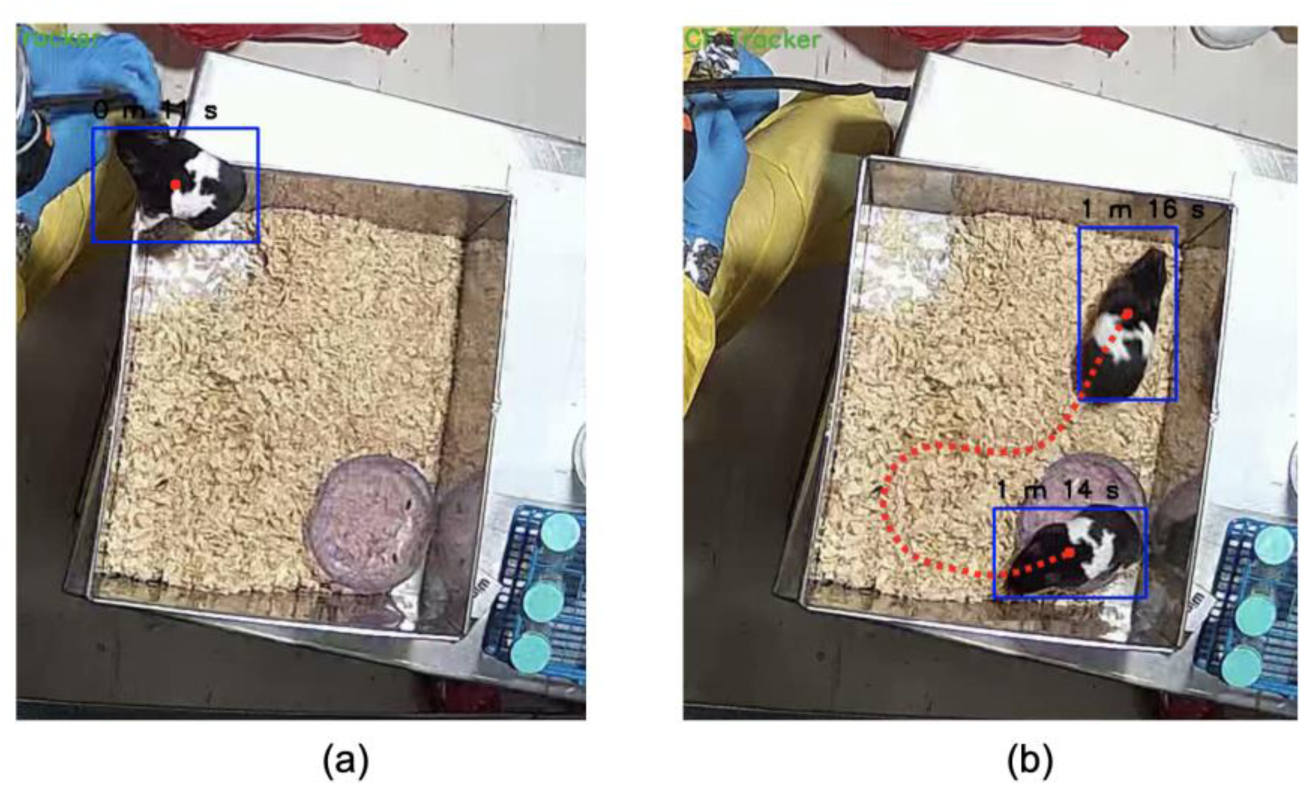
Tracking of animals for body motion description. (**A**) Time intervals that were not considered, where the hamster was on the edges. (**B**) Time intervals considered, where the hamster remained outside the edges and had free movement.

